# Long-term multi-meta-omics resolves the ecophysiological controls of seasonal N_2_O emissions

**DOI:** 10.1101/2024.04.17.589950

**Authors:** Nina Roothans, Martin Pabst, van Diemen Menno, Claudia Herrera Mexicano, Marcel Zandvoort, Thomas Abeel, van Loosdrecht Mark, Michele Laureni

**Affiliations:** Delft University of Technology, Mekelweg 5, 2628 CD Delft, the Netherlands; Waternet, Korte Ouderkerkerdijk 7, P.O. Box 94370, 1090 GJ Amsterdam, the Netherlands; Broad Institute of MIT and Harvard, 415 Main Street, Cambridge, MA 02142, United States of America; Center for Microbial Communities, Department of Chemistry and Bioscience, Aalborg University, Aalborg, Denmark

**Keywords:** microbial ecophysiology, microbial communities, nitrous oxide, wastewater treatment, multi-meta-omics

## Abstract

The potent greenhouse gas nitrous oxide (N_2_O) originates primarily from natural and engineered microbiomes. Emission seasonality is widely reported while the underlying metabolic controls remain largely unresolved, hindering effective mitigation. We use biological wastewater treatment as tractable model ecosystem over nearly two years. Long-term metagenomic-resolved metaproteomics is combined with *ex situ* kinetic and full-scale operational characterization. By leveraging the evidence independently obtained at multiple ecophysiological levels, from individual genetic potential to actual metabolism and emergent community phenotype, the cascade of environmental and operational triggers driving N_2_O emissions is resolved. We explain the dynamics in nitrite accumulation with the kinetic unbalance between ammonia and nitrite oxidisers, and identify nitrifier denitrification as the prime N_2_O-producing pathway. The dissolved O_2_ emerged as the key actionable parameter for emission control. This work exemplifies the yet-to-be-realized potential of multi-meta-omics approaches for the mechanistic understanding and ecological engineering of microbiomes, ultimately advancing sustainable biotechnological developments.

## Introduction

The yearly anthropogenic emissions of nitrous oxide (N_2_O), currently the third most important greenhouse gas, are projected to increase by 50% in the coming 50 years if no mitigation strategies are employed ^1^. N_2_O is mainly produced by microbial communities in natural, managed and engineered ecosystems ^2^. Yet, the mechanisms governing biological N_2_O emissions remain largely unknown. The main challenge lies in the coexistence of nitrogen-converting guilds in complex microbiomes, each emitting N_2_O under a range of complementary conditions that alternate or overlap in most ecosystems (e.g. alternating oxic-anoxic conditions in wastewater treatment plants ^3^ and sea sediments ^4^; substrate concentration gradients in oceans ^5^, soils ^6^ and wastewater treatment biofilms ^7^). In general, high ammonium (NH_4_^+^) and oxygen (O_2_) concentrations stimulate N_2_O production through hydroxylamine (NH_2_OH) oxidation by ammonia-oxidising bacteria (AOB), while high nitrite (NO_2_^−^) and low O_2_ concentrations enhance the nitrifier denitrification pathway ^8^ (Fig. 1A). High NO_2_^−^ and high O_2_ concentrations result in N_2_O accumulation from imbalanced denitrification by heterotrophic denitrifying bacteria (DEN) ^8^ (Fig. 1A). Seemingly ubiquitous is the strong seasonality of N_2_O emissions in many natural and managed environments, such as oceans ^9,10^, soils ^11–13^, lakes ^14,15^ and rivers ^16^, and engineered systems such as wastewater treatment plants ^17–24^ (WWTPs, summarized in Table S1). This indicates that seasonally-impacted macroscopic factors directly influence biological N_2_O turnover. Yet, studying the interactions between environmental conditions, complex microbiome dynamics and N_2_O emissions, and capturing the underlying ecological principles is inherently challenging. To this end, we use biological wastewater treatment as a more tractable model ecosystem, as the N_2_O seasonality is well-represented, while other variables (e.g. aeration, biomass concentration) are controlled or extensively monitored ^25^.

**Figure 1.**
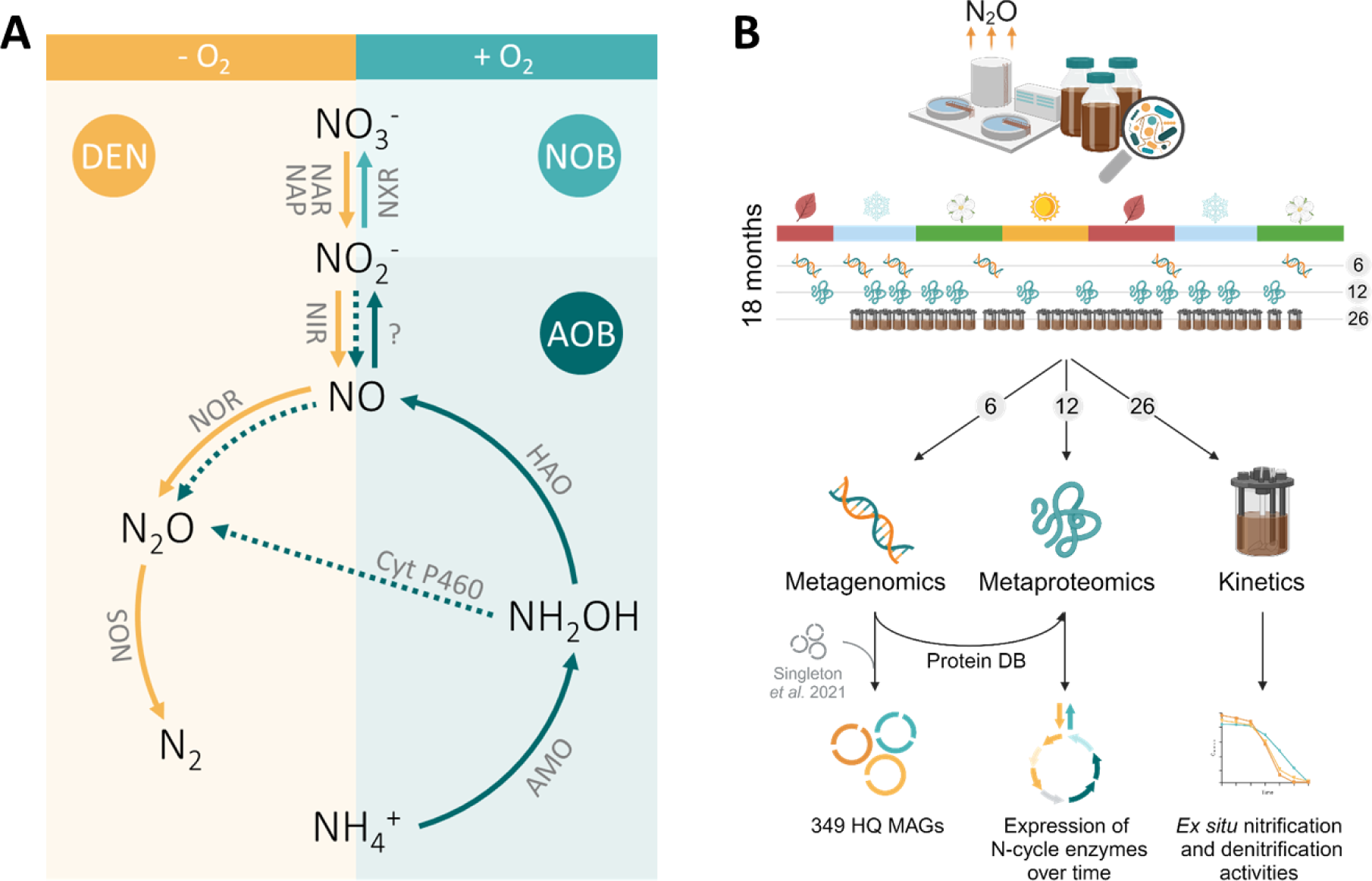
Schematic representation of the nitrogen cycle, experimental approach and obtained datasets. **(A)** Nitrogen conversions in the biological nitrogen removal process and respective enzyme complexes. Ammonia-oxidising bacteria (AOB) aerobically oxidise ammonium (NH ^+^) to hydroxylamine (NH_2_OH) with the ammonium monooxygenase (AMO), NH_2_OH to nitric oxide (NO) with the hydroxylamine oxidoreductase (HAO), and NO to nitrite (NO_2_^−^) with a yet unknown enzyme. AOB can biologically produce N_2_O through the oxidation of NH_2_OH with cytochrome P460 (cyt P460) or through the reduction of NO – produced from NH_2_OH oxidation or nitrifier denitrification (NO ^−^ reduction with the nitrite reductase NIR) – with the nitric oxide reductase (NOR) (dotted arrows). Nitrite-oxidising bacteria (NOB) aerobically oxidise NO ^−^ to nitrate (NO ^−^) with the nitrite oxidoreductase (NXR). Normally under anoxic conditions, denitrifying bacteria (DEN) reduce NO ^−^ to NO_2_^−^ with the membrane-bound or periplasmic nitrate reductase (NAR, NAP), NO ^−^ to NO with NIR, NO to N_2_O with NOR and N_2_O to N_2_ with the nitrous oxide reductase (NOS). Some DEN perform only some steps of the denitrification pathway while others perform the entire pathway. **(B)** Overview of the methodological approach adopted in this study for the eighteen-months characterization of a full-scale WWTP to resolve the microbial mechanisms underlying seasonal N_2_O emissions. Sludge samples were used for metagenomics (6 samples), metaproteomics (12 samples) and *ex situ* activity tests at 20 °C (26 samples). (Created with BioRender.com.)

Most WWTPs emit the majority of their yearly N_2_O during a winter or spring peak lasting 3-4 months, with simultaneous NO_2_^−^ accumulation ^17,21–24,26^ (Table S1). Similarly, higher N_2_O emissions during colder seasons are widely reported for oceans ^10^, soils ^12,13^, and lakes ^14^. Low or increasing temperatures have been hypothesized as the underlying causes for the seasonal N_2_O emissions, but a clear correlation is often missing ^10,13,14,18,19,27,28^. The immediate effect of diverse environmental and process parameters on the N_2_O production rates of AOB and DEN largely explain the short-term N_2_O dynamics in WWTPs ^3,29^ and natural environments ^5,6,30,31^, but fail to describe the widely observed seasonality. Emblematic is the reported higher N_2_O production by AOB at high temperatures ^32^, while most seasonal emissions occur in winter. Broadly applied correlation analyses between N_2_O and environmental and operational parameters have proved insufficient to explain seasonal emissions in WWTPs ^18,24,33^, oceans ^9,10^, soils ^11–13^ and freshwater systems ^14–16^. Despite the evident central microbial role in N_2_O conversions, most studies do not take potential seasonal dynamics of the microbiome’s metabolism into account, likely overlooking key mechanisms linking environmental triggers and emissions. A delay between triggers, metabolic adaptations and emergent phenotype is expected in slow-growing natural and WWTP communities ^28^. Only few studies investigated microbial dynamics during seasonal nitrogen oxides peaks in WWTPs with seemingly contradicting results. Seasonal NO_2_^−^ and N_2_O accumulation events have been attributed to decreased nitrite-oxidising bacteria (NOB) 16S rRNA gene abundances ^19,23^ and increased difference between AOB and NOB activity ^17,22^, while in other instances no seasonal fluctuations were observed in the nitrifying community ^34^. To date, the operational and metabolic mechanisms controlling seasonal N_2_O emissions remain largely unknown, hindering effective mitigation.

We combine long-term metagenomic-resolved metaproteomic analyses with *ex situ* kinetic and full-scale process characterizations to address the mechanistic gap in seasonal N_2_O emissions. The cascade of environmental and operational triggers underlying N_2_O emissions is resolved by leveraging the evidence obtained at multiple ecophysiological levels, from individual genetic potential to actual metabolism and emergent community phenotype. We identify nitrifier denitrification as the prime N_2_O-producing pathway, and the dissolved O_2_ as the central operational parameter to minimize emissions. This work exemplifies the yet-to-be-realized potential of multi-meta-omics approaches to inform ecologically-driven strategies for the management and engineering of microbiomes, ultimately advancing sustainable biotechnological developments.

## Results

### Signature metabolite accumulation profiles

The ecophysiological response of N_2_O-emitting complex microbial communities to seasonal environmental and operational dynamics was studied using the Amsterdam-West wastewater treatment plant (WWTP) as model ecosystem (Fig. 1A-B). The monitoring and sampling period lasted eighteen months and covered two highly comparable N_2_O emission peaks (Fig. 2). The peaks occurred during periods with low water temperatures, namely Feb – May 2021 and Nov 2021 – Mar 2022, and were preceded by the sequential accumulation of NH_4_^+^, O_2_, and NO_2_^−^ (Figs. 2 and S2). The same trend was followed in the five years prior to this study (data not shown). Central to the plant operation is the control of the dissolved O_2_ (DO) concentration as a function of the residual NH_4_^+^ concentration in the aerated compartment. To counteract the temperature-induced nitrification rate reduction, and consequent NH_4_^+^ concentration increase, the weekly average DO concentration was increased from 1 up to almost 3 mg O_2_·L (Fig. 2). In spite of this, O_2_ remained the rate-limiting substrate for nitrification during low temperature periods with high N_2_O emissions, as evidenced by a lower O_2_/NH_4_^+^ ratio in the aerated compartment compared to warmer periods with low N_2_O (Fig. S3). Following the increase in DO, the average NO_2_^−^ concentration in the pooled effluent rapidly increased up to 1.1 mg N·L^−1^. Finally, N_2_O started to accumulate, reaching maximum daily rates of 110 (1^st^ peak) and 101 kg N·d^−1^ (2^nd^ peak) (Figs. 2 and S2). The delay between the maximum DO concentration and the maximum N_2_O emission rate ranged between six and seven weeks for both peaks (Fig. 2), consistent with the imposed average sludge retention time of 11-15 days. Statistically, NO_2_^−^ strongly correlated with the O_2_ concentration (Pearson correlation coefficient of 0.8), and N_2_O with NO_2_^−^ (correlation coefficient 0.7), while they only weakly correlated with all other parameters including the temperature (Fig. S4 and Table S2).

**Figure 2.**
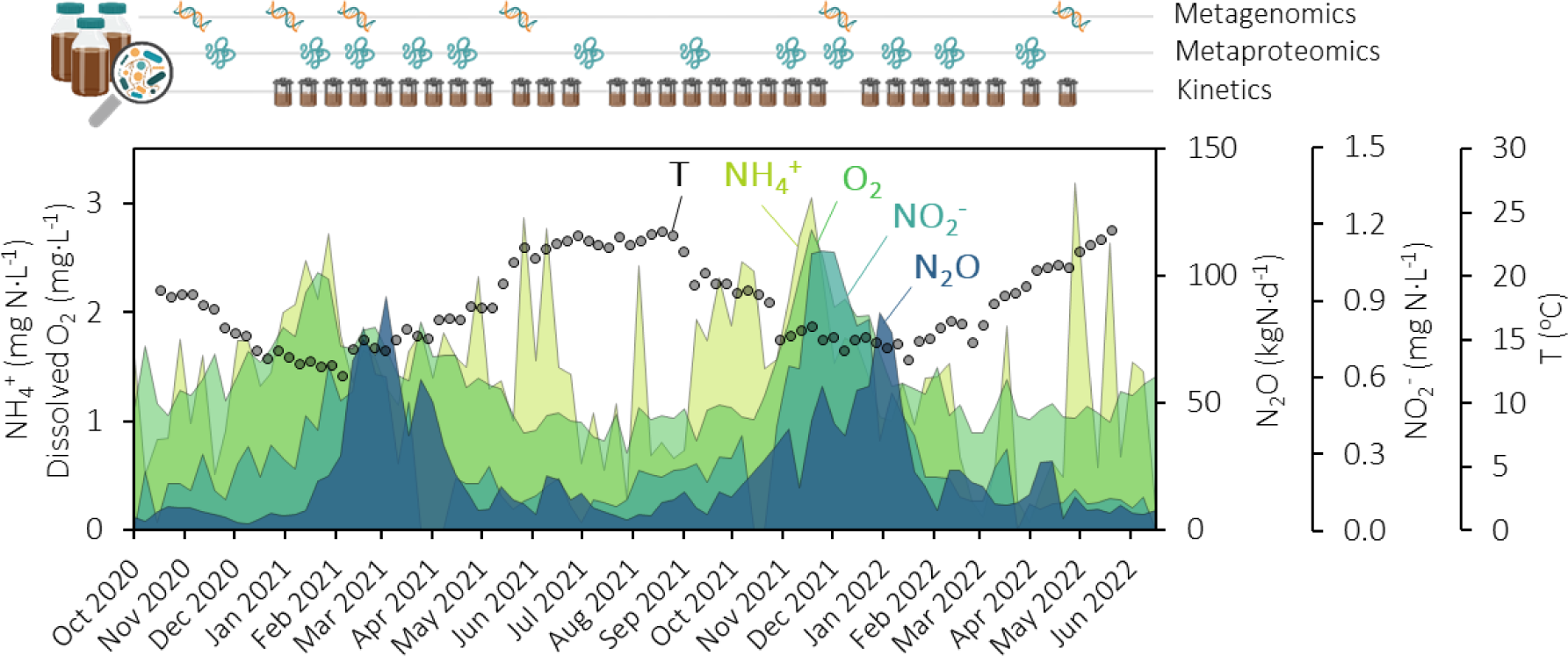
Performance of the wastewater treatment plant (WWTP) monitored during nearly two years (Oct 2020 – Jul 2022) Weekly average parameters at the WWTP, from back to front (light green to dark blue): concentration of NH_4_^+^ and dissolved O_2_ in the nitrification compartment (left axis), pooled effluent NO_2_ ^−^ concentrations (right axis), N_2_O emission rates measured in the off-gas from all reactor compartments (right axis). The water temperature inside the reactor is represented on the right axis (symbols). All metabolites were measured in a single biological nutrient removal lane of the WWTP, except the effluent NO_2_ ^−^ (seven lanes pooled together). Occasional sharp NH_4_ ^+^ peaks were caused by outliers on rainy days (Fig. S2). The scheme above the plot represents the sampling time points for metagenomic (DNA), metaproteomic (protein) and *ex situ* activity tests (bioreactor).

### Maximum nitrogen metabolites conversion rates

To quantify seasonal changes in the microbiome metabolic potential, we estimated every second week the maximum oxidation and reduction rates of the main nitrification (i.e. NH_4_^+^ and NO_2_^−^) and denitrification (i.e. NO_3_^−^, NO_2_^−^ and N_2_O) intermediates, respectively. The maximum NH_4_^+^ oxidation rate almost always exceeded the NO_2_^−^ oxidation rate, with their difference being the highest during the seasonal full-scale metabolite accumulation peaks (Fig. S5). No clear seasonality emerged in the NO_3_^−^, NO_2_^−^, and N_2_O maximum reduction rates, and the N_2_O reduction capacity was 1.4 to 2.1-fold higher than all other nitrifying and denitrifying rates (Fig. S5).

### Genome-resolved taxonomic diversity

The WWTP metagenome was sequenced at six time points to follow the dynamics in microbial composition and functional potential, and to serve as database for the metaproteomic analysis (Fig. 1B). Combined short-read (two samples; average 147 million reads per sample) and long-read DNA sequencing (five samples, one of which also sequenced with short-reads; average 4.3 million reads per sample) resulted in 143 Gbp data, after quality filtering and trimming. A total of 349 high-quality metagenome-assembled genomes (HQ MAGs, ≥ 90% completeness and ≤ 5% contamination) (Fig. 3, Supplementary Data 1) were obtained. The 89 MAGs generated from the five long-read samples were dereplicated with the HQ MAGs from Singleton *et al.* ^35^ at 95% average nucleotide identity of open reading frames to increase the genome-resolved read coverage. From the final 349 HQ MAGs, 44 were unique to our dataset, 268 were unique to the dataset of Singleton *et al.* ^35^, and 37 overlapped between both datasets (Fig. S6). Overall, the HQ MAGs covered 31 phyla and 272 different genera, and included two archaeal species (only bacterial MAGs are represented in Fig. 3). The full 16S rRNA gene was identified in 347 (99.4%) MAGs. The relative abundance of the individual MAGs showed no marked seasonal trend and little variation over the six time points (Fig. S7 and Supplementary Data 1). We therefore discuss the average of their relative abundance among all samples. The two most abundant MAGs belonged to the *Ca.* Microthrix (4.0%) and *Nitrospira* (2.7%) genera (Fig. 3). All other MAGs had an average relative abundance lower than 1%. The majority of the non-nitrifying MAGs contained at least one denitrification gene (DEN, 304) (Fig. 3, Supplementary Data 2). 51 MAGs had the genetic potential to perform dissimilatory nitrite reduction to ammonium (DNRA, containing the *nrfAH* genes), 46 of these also had at least one denitrification gene (Fig. S14, Supplementary Data 2). Seven MAGs harboured the *amoABC* genes (AOB) and eight harboured the *nxrAB* genes (NOB), most of these also had at least one denitrification gene, mainly *nir* and *nor* encoding the NO_2_^−^ and NO reductases, respectively (Fig. S14, Supplementary Data 2). Neither complete ammonia-oxidising (comammox) nor anaerobic ammonia-oxidising (anammox) MAGs were found in the metagenomes.

**Figure 3.**
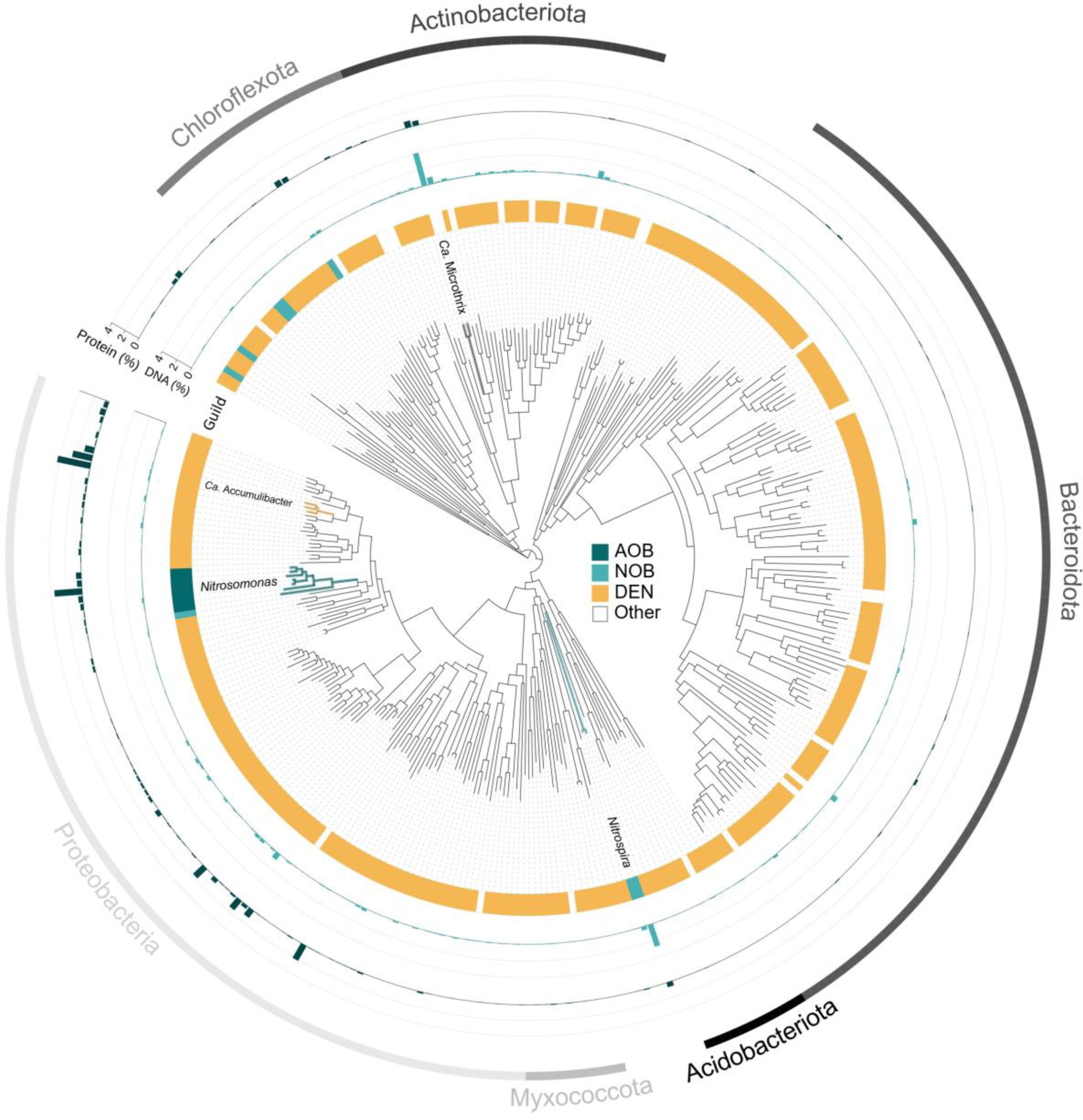
Phylogenetic tree of the 347 bacterial high-quality MAGs extracted from activated sludge (the only two archaeal MAGs are not represented) From the inner to the outer circle: (i) circular phylogenetic tree with the identification of key activated sludge genera *Nitrosomonas*, *Nitrospira*, *Ca.* Accumulibacter and *Ca.* Microthrix; (ii) identification of ammonia-oxidising bacteria (AOB, containing *amoABC* genes, dark blue), nitrite-oxidising bacteria (NOB, containing *nxrAB* genes, light blue) and denitrifying organisms (DEN, non-AOB and non-NOB MAGs harbouring at least one denitrification gene, yellow). Some of the AOB and NOB MAGs also contained one or more denitrification genes (Supplementary Data 2); (iii) average DNA relative abundance of each MAG in the community; (iv) average protein relative abundance of each MAG in the community; (v) identification of the six most abundant phyla.

### Metaproteomic-based functional profile

The dynamics in protein expression of the entire microbial community across twelve samples was assessed by shotgun metaproteomics. We used the protein expression as proxy for active metabolisms and to estimate the protein-based relative abundance of each MAG. In total, 3868 unique protein groups were detected, and 1884 had at least two unique peptides (accounting for 44 ± 1% of the total mass-normalized spectral counts). 1105 of the identified proteins (accounting for 68 ± 1% of the two unique peptides filtered normalized spectral counts) uniquely matched with a single protein predicted in the metagenome (including all MAGs and unbinned sequences). The remaining 779 proteins (accounting for 32 ± 1% of the two unique peptides filtered normalized spectral counts) matched multiple highly similar proteins and could not be linked to a single MAG, yet could be functionally and taxonomically annotated at the genus level. Out of the 349 HQ MAGs, proteins from 143 MAGs (101 genera) were detected (Supplementary Data 1). The HQ MAGs covered 39 ± 1 % of the total protein pool, higher than the 28 ± 4 % coverage of the total community DNA (Fig. 4A). On average, the relative abundance of key activated sludge taxa (e.g. *Ca.* Microthrix, *Ca.* Accumulibacter, *Nitrosomonas* and *Nitrospira*,) differed up to 20-fold between the metagenomic and metaproteomic approaches (Fig. S12). For example, the AOB:NOB ratio was 0.1 in the metagenome and 3.6 in the metaproteome (discussed in Supplementary Section 6). Taxonomically, the diversity was greatest within the DEN guild (proteins from 124 MAGs were detected) with no clear dominant MAG (Fig. 4B). Owing to this high diversity, many DEN organisms were present in too low abundance to be recovered as MAGs even at the already high sequencing depth employed here (20-25 Gbp per sample). Consequently, DNA sequences from many DEN remained in the unbinned portion of the metagenomes, resulting in the majority of the detected denitrification enzymes, namely nitrate, nitrite and nitrous oxide reductases being assigned to the unbinned fraction (Fig. S15). Proteins from all seven AOB and four NOB MAGs were detected in the metaproteome. The AOB consisted entirely of *Nitrosomonas* MAGs, and were dominated by one MAG (Fig. 4B). NOB were dominated by a *Nitrospira* and a Chloroflexota MAG belonging to the Promineofilaceae family (Fig. 4B), but the alpha- and beta-subunits of the nitrite oxidoreductase (NxrA and NxrB) were only expressed by *Nitrospira* and *Ca.* Nitrotoga (Fig. S15). Almost all detected nitrifying enzymes belonged entirely to the recovered MAGs, highlighting the nearly full coverage of the active nitrifying community by the MAGs (Fig. S15). Throughout the monitoring period, the relative proteomic abundance of DEN hardly fluctuated, and the AOB and NOB guilds fluctuated similarly over time (Fig. 4C). The maximum guild-specific fold change in the proteome was 1.1 (DEN), 1.8 (AOB) and 2.5 (NOB). Overall, there were no major shifts in the MAG-based composition of each guild, at both DNA and protein level (Figs. S8-S10), and there were no significant correlations between protein-level taxa abundance and WWTP performance (Table S4).

**Figure 4.**
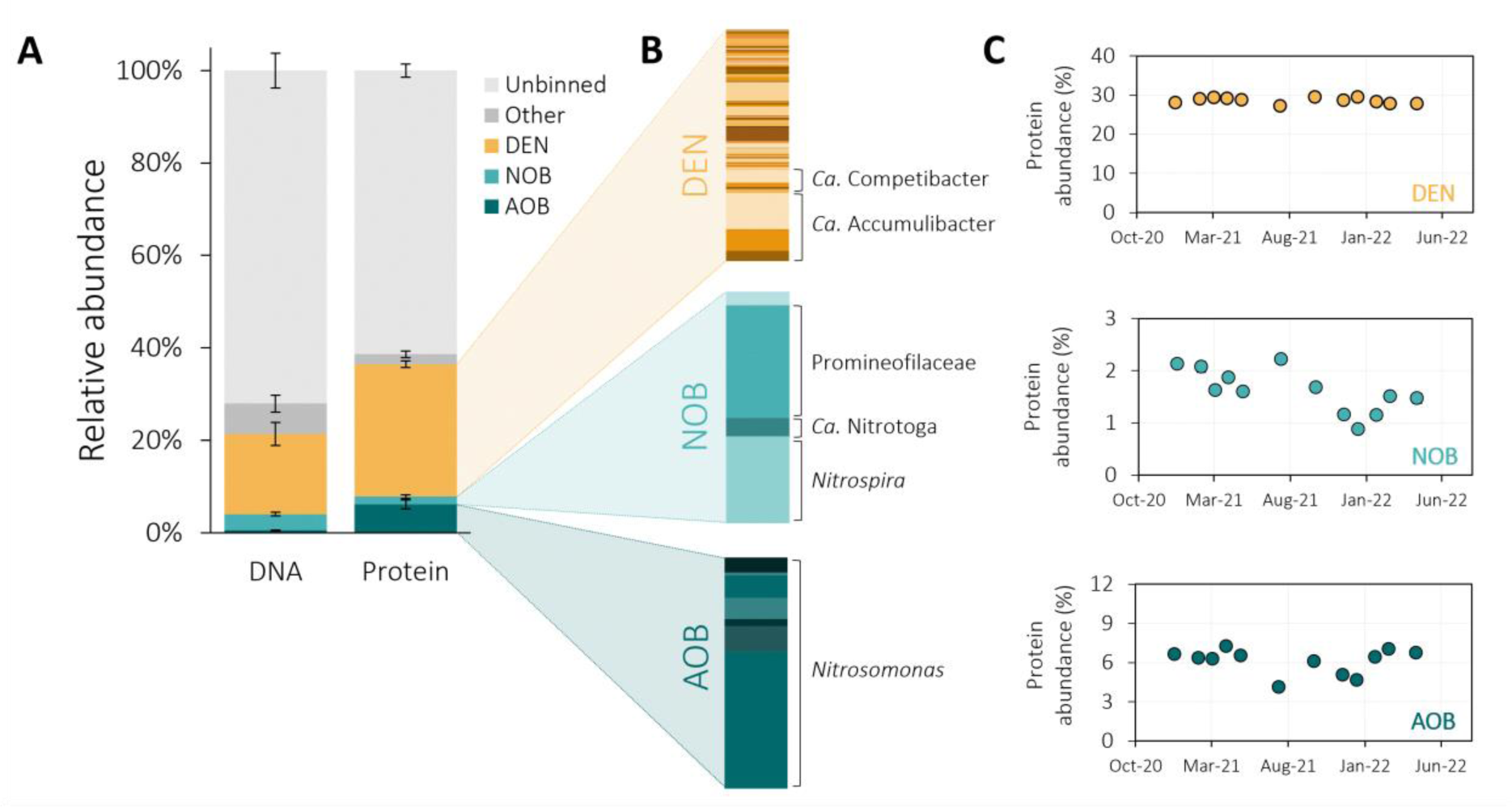
MAG-based functional guild distribution in the metagenomes and metaproteomes of the activated sludge. **(A)** Average relative abundance of denitrifying bacteria (DEN, non-AOB and -NOB MAGs containing at least one denitrification gene, yellow), nitrite-oxidising bacteria (NOB, containing *nxrAB* genes, light blue), ammonia-oxidising bacteria (AOB, containing *amoABC* genes, dark blue), other metagenome-assembled genomes (dark grey) and unbinned sequences (light grey) in the total metagenome (DNA) and metaproteome (Protein) of the activated sludge. Some of the AOB and NOB MAGs also contained one or more denitrification genes (Supplementary Data 2). The error bars represent fluctuations within six (DNA) and twelve (protein) activated sludge samples taken throughout eighteen months. **(B)** MAG-based composition of the DEN, NOB and AOB guilds. The most abundant genera in the DEN (*Ca.* Accumulibacter and *Ca.* Competibacter), NOB (unidentified Promineofilaceae genus, *Ca.* Nitrotoga, Nitrospira) and AOB (*Nitrosomonas*) guilds are highlighted. **(C)** Temporal fluctuations in the relative protein abundance of the DEN (yellow), NOB (light blue) and AOB (dark blue) guilds. The error bars represent standard deviations between technical duplicates and are all smaller than the symbols.

### Unbalanced nitrification drives seasonal nitrite accumulation

The net accumulation and potential emission of any nitrogen intermediate results from the unbalance between its production and consumption rates. Nitrite, a central metabolite exchanged between AOB, NOB and DEN (Fig. 1A), always accumulated prior to the N_2_O peaks (Fig. 2). To understand the NO_2_^−^ flux balance dynamics, we focused on the DNA, expressed proteins and *ex situ* activity ratios of NO_2_^−^-producing and -consuming guilds. At all levels (genomic, proteomic and kinetic), the DEN guild did not display significant seasonal dynamics (Figs. 4C, S8 and S19). Contrastingly, the (un)balance between AOB (NO_2_^−^ producer) and NOB (NO_2_^−^ consumer) fluctuated the most during the monitored period. The ratio between the total abundances of AOB and NOB, both at DNA and protein level, was up to 3-fold higher during periods of high effluent NO_2_^−^ concentrations, compared to the rest of the year (Fig. 5A-B). At individual protein level, including MAG and unbinned proteins, the ratio between the expression of the key NH_4_^+^-consuming enzyme (represented by the beta-subunit of the ammonia monooxygenase – AmoB) and NO_2_^−^-producing enzyme (represented by the hydroxylamine oxidoreductase – Hao) of AOB relative to the catalytic subunit of the NO_2_^−^ oxidoreductase of NOB (NxrA) were also higher (Fig. 5C, Supplementary Data 3). Consistently, the ratio between the maximum NH_4_^+^ and NO_2_^−^ oxidation activities was larger during high NO_2_^−^ concentration periods (Fig. 5D).

**Figure 5.**
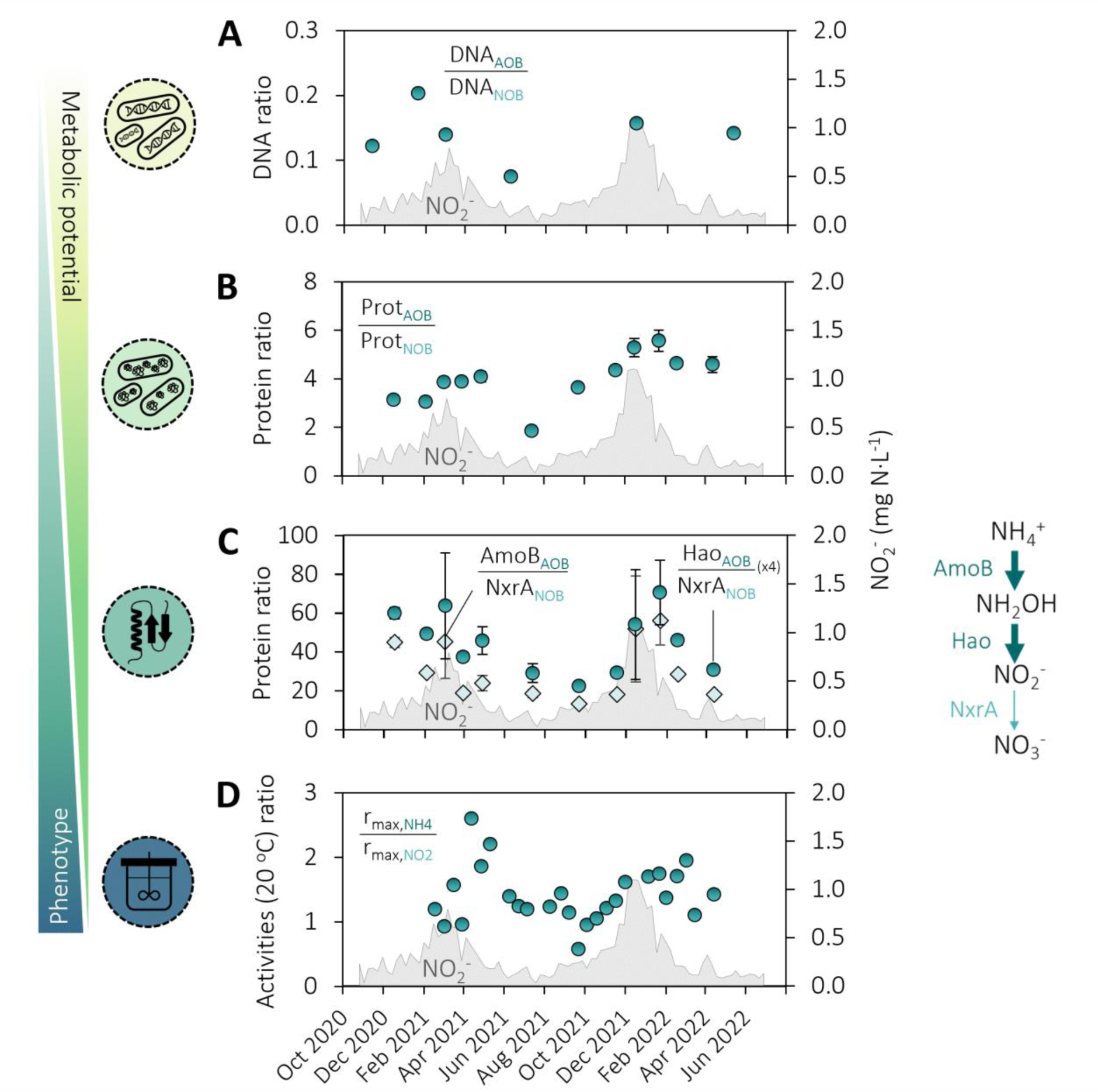
Genomic, proteomic and maximum activity fluctuations of AOB and NOB in activated sludge during periods of high and low nitrite accumulation. **Left axes: (A)** Ratio between the total relative DNA abundance of ammonia-(AOB) and nitrite-oxidising bacteria NOB (circles). **(B)** Ratio between the total relative protein abundance of AOB and NOB (circles). **(C)** Ratios between the relative abundance of NO_2_ ^−^-producing and -consuming enzymes of AOB and NOB, respectively: beta-subunit of the ammonia monooxygenase (AmoB) divided by the catalytic subunit of nitrite oxidoreductase (NxrA) (diamonds); and hydroxylamine oxidoreductase (Hao) divided by NxrA (x4, circles). The enzyme abundances include the proteins belonging to the MAGs and the unbinned fraction. The error bars in all protein ratios were propagated from standard deviations of technical duplicates and some are smaller than the symbols. The respective enzymatic conversions are represented on the right. **(D)** Ratio between the maximum *ex situ* NH_4_ ^+^ and NO_2_ ^−^ oxidation rates measured at 20 °C (circles). **Right axes: (A-D)** Weekly average NO_2_ ^−^ concentration in the effluent (seven parallel lanes pooled together, grey area).

### Overexpressed nitrifier denitrification during N_2_O accumulation

In analogy to nitrite, we used ratios between the relative abundance of enzymes directly or indirectly producing and consuming N_2_O as proxy for the N_2_O flux balance. The total enzyme abundances include MAG and unbinned protein abundances (Supplementary Data 3). The seasonally accumulated NO_2_^−^ can be reduced to N_2_O by both AOB and DEN, sequentially using the Cu-(NirK) or *cd1*-type (NirS) NO_2_^−^ reductases and the nitric oxide reductase (Fig. 1A). Here, NirK and NirS were exclusively expressed by nitrifiers and DEN, respectively (Fig. S15). Four *Nitrosomonas* (AOB) and one *Nitrospira* MAG (NOB) accounted for most of the NirK expression (75% and 17%, respectively) (Fig. S15). Within the nitrifying community, the relative abundance of NirK over the key AOB enzymes AmoB and Hao was the highest during periods of high NO_2_^−^ and N_2_O accumulation (Fig. 6A). The ratio of total relative abundance of NirK over the competing NO_2_^−^-oxidising NxrA (NOB) and NO_2_^−^-reducing NirS (DEN) followed a similar trend (Fig. 6B). NosZ is the only known N_2_O-reducing enzyme, and the ratio NirK/NosZ clearly reflected the seasonal dynamics, being higher during seasonal peaks (Fig. 6C). Similarly, yet to a significantly lower extent, also the ratio between the hydroxylamine (NH_2_OH) producing AmoB and consuming Hao and CytP_460_ (Fig. S18), and the ratio NirS/NosZ (Fig. S19C) displayed some seasonality. The here employed protein extraction protocol does not allow for the quantification of membrane-bound proteins, such as the nitric oxide reductases, which were therefore not included in the discussion.

**Figure 6.**
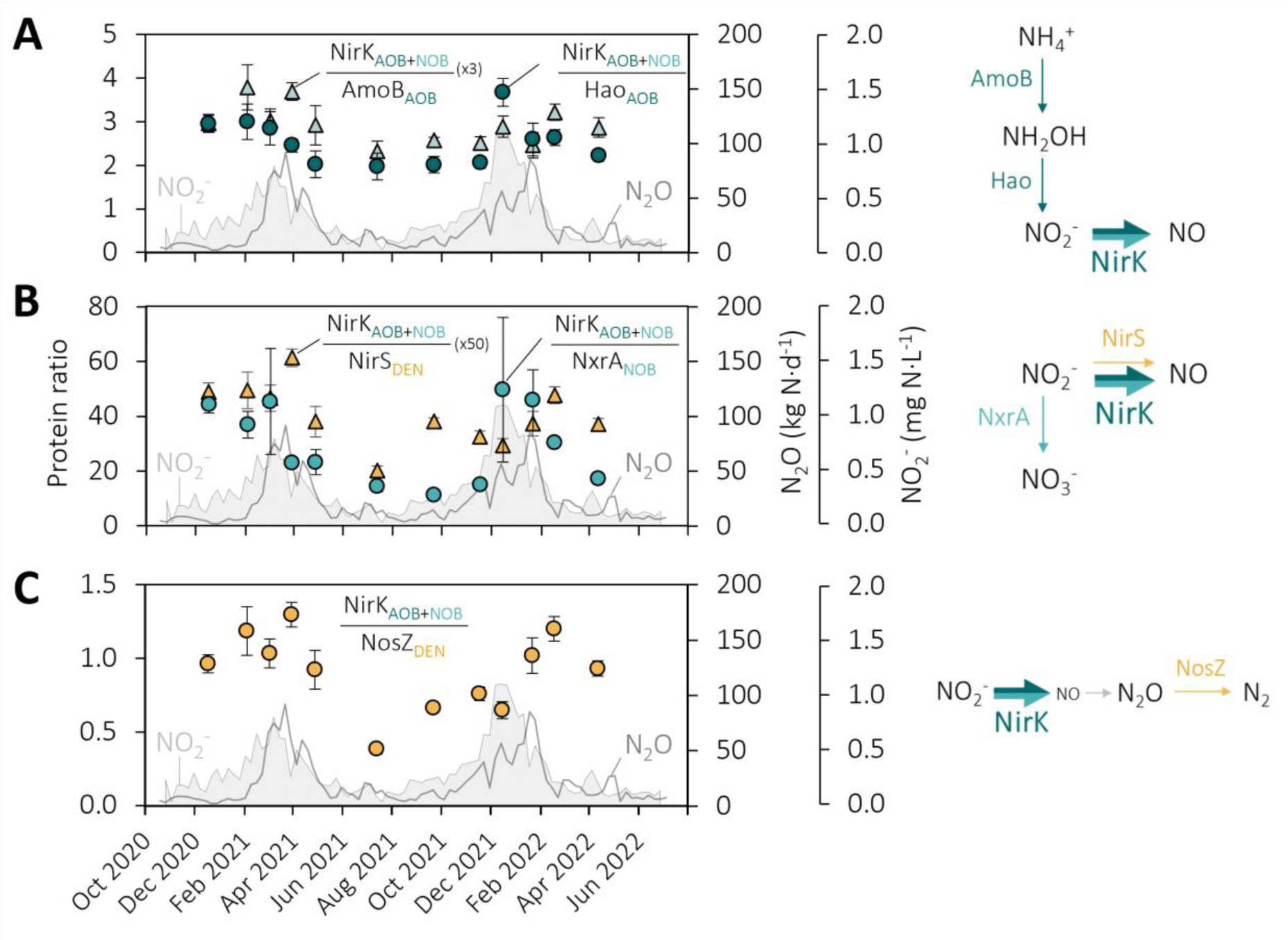
NirK overexpression relative to other nitrogen enzymes during periods of high NO_2_^−^ concentrations and N_2_O emissions. Left axes (symbols) **(A) NirK vs. other AOB enzymes** Ratio between the total relative abundance of NO_2_^−^-consuming NirK and the other key AOB enzymes Hao (circles) and AmoB (triangles). **(B) NirK vs. competing NO_2_^−^ consuming enzymes.** Ratio between the total relative abundance of NO_2_^−^-consuming NirK and the NO_2_^−^ competing NxrA (circles, NOB) and NirS (x50, triangles, DEN). **(C) NirK in N_2_O balance.** Ratio between the total relative abundance of NirK (producing the N_2_O precursor NO) and the only known enzymatic N_2_O-sink N_2_O reductase (NosZ) (circles). The enzyme abundances include the proteins belonging to the MAGs and the unbinned fraction. The error bars in the protein ratios were propagated from standard deviations of technical duplicates. All enzymatic conversions are schematically represented on the right. NirK is expressed by both AOB and NOB, but the activity and function of the enzyme in NOB are yet unknown. **Right axes: (A-C)** Weekly average NO_2_^−^ concentration in the effluent of the WWTP (seven parallel lanes pooled together, grey area) and N_2_O emission rates measured in the off-gas from all the reactor compartments in one lane at the WWTP (grey line).

## Discussion

We postulate that the seasonal accumulation of NO_2_^−^ and subsequent emissions of the potent greenhouse gas N_2_O at a full-scale WWTP are related to fluctuations in the balance of key nitrogen-converting populations, rather than their individual abundance or activity. No major changes in the DNA and protein composition, nor significant correlations with plant performance, were observed throughout eighteen months of operation. This is consistent with previous metagenomic and 16S rRNA gene amplicon sequencing reports in WWTPs ^36–39^. The microbiome was dominated by a taxonomically diverse DEN community (74% of the binned community proteome), in line with most genomic and transcriptional analyses of conventional WWTPs ^19,40,41^. While the high DEN abundance may have masked fluctuations at guild level, the absence of significant changes at the activity and individual protein level further supports the DEN stability. Instead, the DNA and protein abundances of the nitrifying community, dominated by one AOB and two NOB MAGs, fluctuated over time, yet not consistently with the observed nitrogen oxides accumulation dynamics. This aligns with most studies reporting limited to no correlation between AOB and NOB 16S rRNA gene abundances and seasonal nitrification failures ^34^, or AOB and NOB conversion rates and N_2_O production ^22^. Only few studies observed a correlation between increased N_2_O emissions and increased relative AOB abundances (16S) ^42^, AOB *ex situ* activities ^43^, or decreased NOB abundances (16S) ^19,23^. Yet evidence remains sparce and seemingly conflicting, ultimately hindering mechanistic generalizations. This lack of general consensus resides in the fundamental dependency between metabolite dynamics and the trade-off between their production and consumption rates (i.e. the balance between the producing and consuming guilds), rather than their individual magnitudes.

Against a relatively stable DEN community, featuring a fairly constant nitrite production and reduction potential, we identified the unbalance between AOB (NO_2_^−^ producer) and NOB (NO_2_^−^ consumer) as the primary cause for seasonal nitrite accumulation. During the nitrite peaks preceding the N_2_O ones, a higher ratio of AOB over NOB was observed at genomic, proteomic and kinetic levels. To date, only Bae et al.^22^ quantitatively linked N_2_O emissions with increased AOB/NOB *ex situ* activity ratios in an otherwise stable nitrifying community based on 16S rRNA gene sequencing. Gruber et al.^23^ observed stable AOB but lower NOB and filamentous bacteria 16S rRNA gene abundances during winter N_2_O emissions, and hypothesized a selective NOB washout due to compromised floc integrity. Here, the fluctuations in the sludge settleability (representing floc integrity) and in the DNA and protein abundances of *Ca.* Microthrix (filamentous bacteria) did not follow the full-scale metabolite profiles, nor the NOB abundance or the AOB/NOB ratio (Figs. S2, S11 and Table S2). The known higher sensitivity of NOB to the toxic free ammonia and nitrous acid compared to AOB ^44–46^ has also been suggested as potential cause for nitrite accumulation ^45^. However, in our case, the estimated concentration of free ammonia (0.03 mg N·L^−1^) and nitrous acid (0.001 mg N·L^−1^) were far below the NOB toxicity thresholds (Tables S6-S7) ^44–47^. Instead, we argue that the unbalanced AOB/NOB ratio results from a cascade of separate environmental and operational perturbations differentially impacting their respective growth rates (Fig. 7). The decrease in temperature reduces both AOB and NOB growth rates, and may alone promote the selective washout of the slower-growing NOB (as estimated in this work and consistent with literature values; Table S8, Fig. S21). In addition, reduced AOB rates lead to the accumulation of ammonium, with the operationally undesired worsening of effluent quality. In response, most WWTPs increase the operational dissolved O_2_ set point to promote nitrification. The increased availability of ammonium selectively favours AOB, while, in principle, the increase in dissolved O_2_ positively impacts the growth rate of both AOB and NOB. However, the reported lower AOB apparent affinity for O_2_ in activated sludge ^48–51^ likely favours AOB over NOB, further enhancing the initial differential temperature impact on their growth rates. Ultimately, nitrite accumulation is the result of the progressive relative enrichment of AOB over NOB. To test our hypothesis, we developed and implemented a simple mathematical model based on the experimentally estimated kinetic parameters and literature-derived stoichiometric parameters (Tables S9-S12). The model reproduced all observed seasonal metabolites peaks induced by decreasing temperatures and consequent increase in ammonium and operational dissolved O_2_. The simulations also captured the progressive relative biomass increase of AOB over NOB (Fig. S23). These results strongly indicate that the sequential seasonal nitrogen oxides peaks result from a cascade of distinguishable events, where temperature is the initial trigger but not the sole direct cause, as commonly hypothesized. The absence of a single parameter correlating with nitrite and subsequent N_2_O emissions likely explains the difficulties of past studies to identify direct correlations ^18,24,33^. Importantly, the dissolved O_2_ concentration emerged as the central operational parameter to act upon, and we posit that the AOB/NOB unbalance may be largely prevented by anticipating in time, i.e. before measurable NH_4_^+^ accumulation, the operational O_2_ increase.

**Figure 7.**
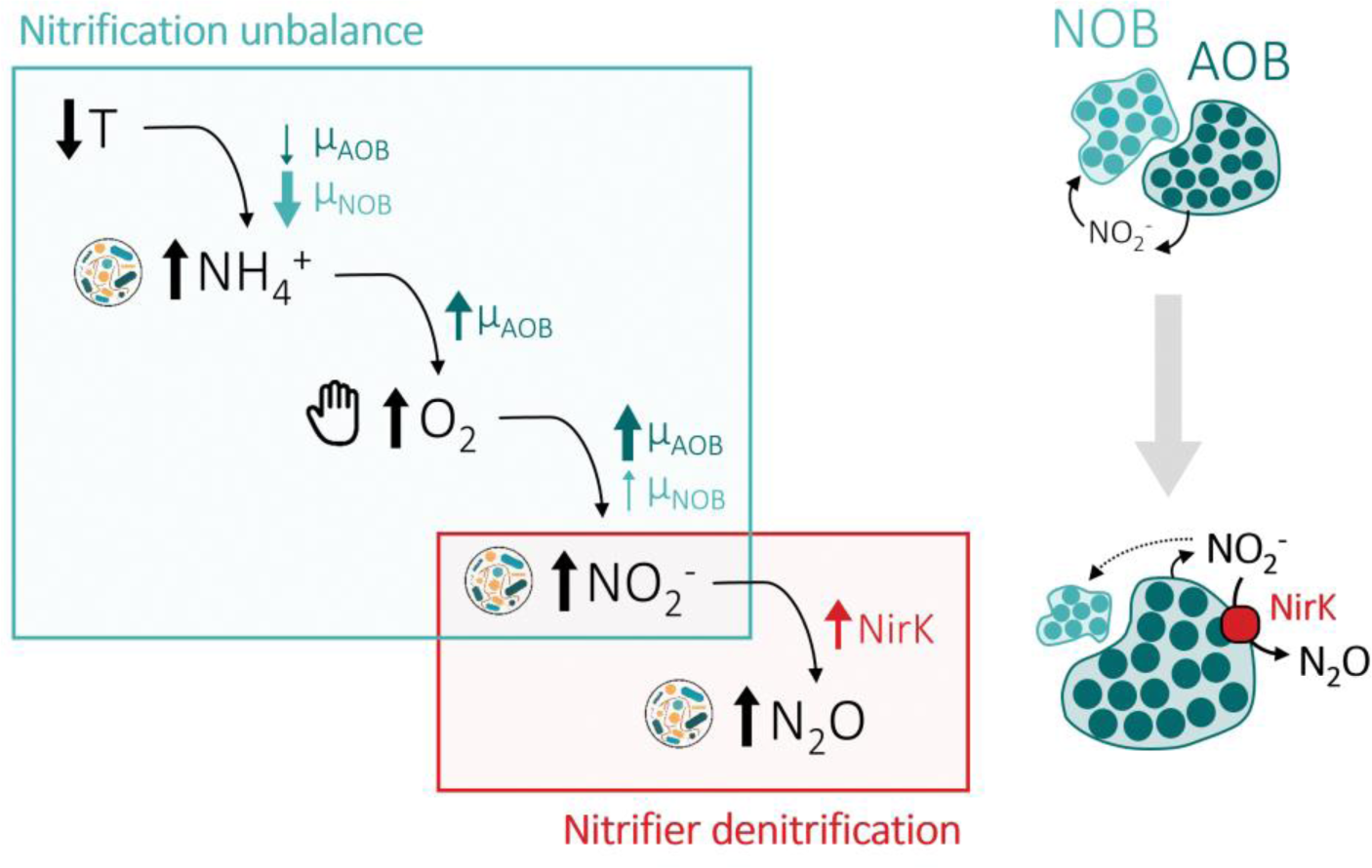
Schematic representation of the proposed ecophysiological cascade underlying seasonal N_2_O emissions in WWTPs. A decrease in temperature causes lower growth rates of ammonia-(AOB) and nitrite-oxidising bacteria (NOB), promoting ammonium accumulation and a selective washout of the slower-growing NOB; the resulting increased ammonium concentrations stimulate the growth of AOB and induce the process control to increase the operational dissolved O_2_ concentration; the increased O_2_ concentrations increase the growth rates of both AOB and NOB, but may selectively benefit AOB with a lower apparent affinity for O_2_. The resulting increased AOB/NOB ratio causes the accumulation of nitrite and consequent stimulation of nitrifier denitrification by AOB, as observed in the overexpression of the Cu-type nitrite reductase (NirK). The ammonium, nitrite and N_2_O concentration increases are a result of changes in the microbial community metabolism, while the increase in O_2_ concentration is the only manually controlled parameter in the cascade.

The last metabolite to accumulate along the reconstructed ecophysiology cascade is N_2_O. High nitrite concentrations are well-known to lead to N_2_O emissions through both nitrifier and heterotrophic denitrification ^3^, yet the dominant pathway underlying seasonal N_2_O emissions remains unclear ^21,24,43^. We use the nitrite reductases (NirK and NirS) as proxy for N_2_O production, and their genome-resolved taxonomy to differentiate between nitrifier and heterotrophic denitrification. Considering the fast turnover of NO ^8^, the use of Nir allows to overcome the challenges in detecting the membrane-bound hydrophobic nitric oxide reductase in metaproteomic analyses ^52,53^. Unbalanced heterotrophic denitrification was excluded as the main N_2_O producing pathway during the seasonal emissions owing to the relatively constant ratio between NirS and NosZ, both exclusively expressed by DEN, and their rates. The nitrite reductase NirK was exclusively expressed by nitrifiers, primarily by AOB, so it was used as proxy for nitrifier denitrification. NOB *Nitrospira* contributed to about one fifth of the total detected NirK, but its activity and function remain unknown ^54–56^. A marked increase in the ratio of NirK over other AOB enzymes (AmoB and Hao) and the competing NO_2_^−^-consuming enzymes (NxrA from NOB and NirS from DEN) was observed during the seasonal nitrogen oxide peaks. The higher expression of NirK was likely induced by the seasonally increased nitrite concentrations ^57,58^, and suggests an increased relative nitrite flux towards nitrifier denitrification rather than nitrite oxidation or heterotrophic nitrite reduction. Emissions also coincided with periods in which O_2_ was identified as the metabolically limiting substrate for AOB (i.e. lower O_2_/NH_4_^+^ ratios compared to the rest of the year), likely forcing AOB to resort to nitrifier denitrification as additional electron sink ^59,60^. The observed slight imbalance between hydroxylamine-producing AmoB and -consuming Hao and Cyt P460 makes it here tempting to speculate that hydroxylamine accumulated as a result of the kinetic O_2_ limitation ^59^, further supporting an electron unbalance in the AOB metabolism. To date, only one report suggested a correlation between N_2_O emissions in WWTPs and *nirK* gene transcripts abundance, quantified by RT-qPCR ^61^. Yet, the *nirK* transcripts were not taxonomically classified and were assumed to entirely belong to heterotrophic denitrifiers ^61^. All other studies discussing seasonal N_2_O emissions in WWTPs infer the main N_2_O-producing pathways based on metabolite profiles, and a general consensus is still lacking ^18,20,21,24,43^ (Table S1). Gruber et al. ^62^ suggest heterotrophic denitrification as the main N_2_O-producing pathway in a WWTP using natural isotopic signatures, but seasonal dynamics were not captured. More importantly, the isotopic signatures of N_2_O produced through nitrifier and heterotrophic denitrification largely overlap, challenging the possibility to univocally distinguish the two pathways ^62,63^. For the same reason, ^15^N/^18^O tracer methods also did not yield conclusive results ^63^. Instead, by integrating metagenomic-guided metaproteomics with kinetic analyses and full-scale operational data we provide independent evidence on multiple ecophysiological levels identifying nitrifier denitrification as the prime N_2_O-producing pathway during seasonal emissions. More broadly, our results demonstrate the untapped potential of multi-meta-omics integration in biotechnological developments to resolve the complexity and advance the engineering of the underlying microbiomes.

## Methods

### WWTP operation

The covered Amsterdam-West WWTP has the daily capacity to treat 200,000 m^3^ municipal wastewater under dry weather conditions (1 million population equivalents). After fine screening and primary sedimentation, carbon, phosphorus and nitrogen are biologically removed in a modified University of Cape Town configuration in seven independent parallel cylindric plug-flow activated sludge tanks (Fig. S1). Nutrient removal occurred in four compartments: anaerobic (biological phosphorus removal), anoxic (denitrification), facultative (aerated when additional nitrification capacity was required), and aerobic (nitrification) (Fig. S1). The setpoint for the dissolved O_2_ concentration in the aerobic and facultative zones was set as function of the measured NH_4_^+^ concentration in the aerated compartment. The average sludge retention time (SRT) was 11-15 days and was controlled to maintain an average total suspended solids of 4.2 g·L^−1^. N_2_O was measured in the combined gas exhaust of all compartments (anaerobic + anoxic + facultative + aerobic) of a single lane using an Rosemount^TM^ X-STREAM gas analyser (Emerson). NH_4_^+^, NO_3_^−^ and N_2_O were measured in a single biological nutrient removal lane of the WWTP, NO_2_^−^ was measured in the pooled effluent of seven lanes.

### *Ex situ* batch activity tests with full-scale activated sludge

The maximum nitrification and denitrification activities of the activated sludge were measured every two weeks between January 2021 and May 2022. For consistency, the sludge sampling, handling and storage, and the activity tests were always performed in the same manner. Samples were collected from the aerated compartment of the monitored full-scale activated sludge reactor and stored in two litre glass bottles in the fridge for a maximum of four hours. The sludge was transported under cold conditions (never reaching a temperature above 10 °C) and immediately placed in a 3 L jacketed glass bioreactor with a 2 L working volume (Applikon, Getinge). The sludge was made anoxic by sparging with N_2_ for 1 h at 0.5 L·min^−1^ (after which the bioreactor was sealed) and was incubated overnight with 50 mg N·L^−1^ NaNO_3_ to consume the internal carbon storages. During overnight storage and subsequent activity tests, the sludge was stirred at 750 rpm by two six-blade turbines, the temperature was maintained at 20 ± 1 °C using a cryostat bath (Lauda), and the pH was automatically maintained at 7.0 ± 0.1 by 1 M HCl and 1 M NaOH with two peristaltic pumps (Watson Marlow) controlled by an in-Control process controller (Applikon, Getinge). The pH and dissolved oxygen were continuously monitored with probes (Applikon AppliSens, Getinge). Influent gas flows were controlled by mass-flow controllers (Brooks). After overnight incubation with NO_3_^−^, the sludge was activated by adding a spike of NaNO_3_ (5 mg N·L^−1^) and a mixture of organic carbon (acetate, pyruvate, glucose, 37.5 mg COD·L^−1^ each). The batch activity tests were sequentially performed on the same day in the following order: N_2_O, NO_2_^−^ and NO_3_^−^ reduction (denitrification), and NH_4_^+^ and NO_2_^−^ oxidation (nitrification) (Table 2). Before each batch, the depletion of the previous nitrogen compound was ensured. Substrates were added to the bioreactor with a syringe and needle through a rubber septum, marking the start of the batches. The batches’ progress was monitored with NO_2_^−^ and NO_3_^−^ MQuant^®^ colorimetric test strips (Merck).

Nitrogen compounds were added at 12 mg N·L^−1^, in the form of N_2_O (sparging 1.5% N_2_O + 98.5% N_2_ at 0.5 L·min^−1^ during 15-20 min), NaNO_2_ (1.2 mL), NaNO_3_ (1.2 mL) and NH_4_HCO_3_ (1.2 mL) from concentrated stocks. The proportion bicarbonate to nitrogen was kept the same for the two nitrification batches by supplying 0.9 mM NaHCO_3_ to the NO_2_^−^ oxidation batch. The organic carbon compounds were added at the start of the denitrification batches (each 75 mg COD·L^−1^, at least 2-fold higher than stoichiometrically needed) from anoxic concentrated stock solutions: sodium acetate (C_2_H_3_NaO_2_, 3 mL), sodium pyruvate (C_3_H_3_NaO_3_, 3 mL) and glucose (C_6_H_12_O_6_, 3 mL). The concentration of pyruvate was 4-fold lower in the batch tests from January until mid-August 2021, but this had no effect on the measured activities. Before each denitrification test, anoxic conditions were ensured by sparging N_2_ at 0.5 L·min^−1^ for 20 min, after which the reactor was sealed off from the exterior. The transition from anoxic to oxic conditions was achieved by sparging air at 0.5 L·min^−1^ for at least 1 h. During each nitrification test, oxic conditions (> 70% air saturation) were ensured by continuously sparging air at 0.5 L·min^−1^. When necessary, foam formation was reduced with a few drops of six times diluted antifoam C 391 emulsion (Merck Life Science NV). For supernatant analysis, samples were taken every 3, 5, 10 or 15 min (depending on the length of the batches), and immediately filtered with a 0.45 µm PVDF Millex syringe filter (Merck) and placed on ice. The samples were stored at 4 °C until analysis on the following day.

**Table 1.** Order and details of the nitrification and denitrification activity tests performed on a single day, every second week. The denitrification tests (N_2_O, NO_2_ ^−^ and NO_3_ ^−^ reduction) were performed under anoxic conditions, with a mixture of organic carbon compounds as electron donor. Prior to each denitrification batch the broth was sparged with N_2_ during 20 min to ensure anoxic conditions and remove intermediate nitrogenous gases. The nitrification tests (NH_4_ ^+^ and NO_2_ ^−^ oxidation) were performed with O_2_ as electron acceptor, under continuous aeration. Between the denitrification and nitrification batches, the broth was made oxic by sparging air for 60 min. Each nitrogen compound was added at a final concentration of 12 mg N·L^−1^.

| Batch | Electron donor | Electron acceptor | Length (min) | Sparging | Conditions |
| --- | --- | --- | --- | --- | --- |
| N<sub>2</sub>O reduction (DEN) | Acetate, pyruvate, glucose | N<sub>2</sub>O | 24 - 105 | Off | Anoxic |
| NO<sub>2</sub><sup>-</sup> reduction (DEN) | Acetate, pyruvate, glucose | NO<sub>2</sub><sup>-</sup> | 25 - >150 | Off |  |
| NO<sub>3</sub><sup>-</sup> reduction (DEN) | Acetate, pyruvate, glucose | NO<sub>3</sub><sup>-</sup> | 35 - >150 | Off |  |
| NH<sub>4</sub><sup>+</sup> oxidation (AOB) | NH<sub>4</sub><sup>+</sup> | O<sub>2</sub> | 30 - >150 | Air | Oxic |
| NO<sub>2</sub><sup>-</sup> oxidation (NOB) | NO<sub>2</sub><sup>-</sup> | O<sub>2</sub> | 45 - >150 | Air |  |

### Analytical methods

The concentrations of NH_4_^+^, NO_2_^−^ and NO_3_^−^ in the filtered supernatant were spectrophotometrically measured on the day following the batches, using the Gallery^TM^ Discrete Analyzer (Thermo Fisher Scientific) or cuvette test kits (LCK339, LCK342 and LCK304, Hach Lange). When measuring NO_3_^−^ with the cuvette test kits, the samples were diluted 1:1 with 20 g·L^−1^ sulfamic acid to remove NO_2_^−^ as interference. The volatile suspended solids concentration (ash content subtracted from the dried biomass), measured in triplicate, was taken as proxy for the biomass concentration. Immediately upon arrival, 3x 25 mL of sludge was centrifuged at 4200 rpm for 20 min, the pellet was resuspended in 15 mL MilliQ water, dried at 105 °C (24 h) and burned at 550 °C (2 h). The concentrations of O_2_, CO_2_ and N_2_O in the condenser-dried reactor off-gas were monitored by a Rosemount NGA 2000 off-gas analyser (Emerson). The dissolved N_2_O concentrations were monitored and recorded every minute with a standard N_2_O-R microsensor (customized concentration range 0.4 – 2 mM, Unisense) and a picoammeter PA2000 (Unisense). The dissolved N_2_O concentrations were calculated using the average of all calibrations performed 1-2 days before every batch series.

### Calculations activity tests

The maximum NO_2_^−^ and NO_3_^−^ reduction and NH_4_^+^ and NO_2_^−^ oxidation rates were obtained through linear regression of the substrate concentration profiles over time. The slope was determined using at least four concentration points in the linear range. The maximum N_2_O reduction rate was calculated in Spyder IDE v5.1.5 using Python v3.9.12 and the NumPy v1.21.5 ^64^, SciPy v1.7.3 ^65^ and Pandas v1.4.2 ^66^ packages, taking into account the gas-liquid transfer between the reactor broth and headspace throughout the batch test (Supplementary Section 13). A system of ordinary differential equations (ODEs), representing the liquid and headspace mass balances, was defined to describe the gas-liquid transfer over time:

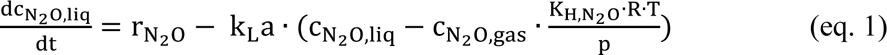

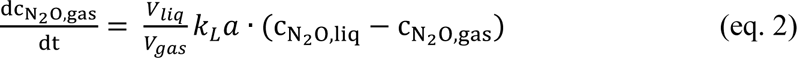

With c_N2O,liq_ and c_N2O,gas_ the N_2_O concentration in the liquid and headspace, r_N2O_ the unknown N_2_O consumption rate, k_L_a the experimentally determined volumetric mass transfer coefficient (5 h^−1^), K_H,N2O_ the Henry coefficient (27.05 mM/atm), R the ideal gas constant (8.206 x 10^−5^ L·atm·K^−1^·mmol), T the temperature, p the pressure, and V_liq_ and V_gas_ the broth and headspace volumes. The rates were obtained by fitting the model to the experimental data, i.e. by minimizing the sum of squared errors between the experimentally measured and calculated (eq. 1-2) N_2_O concentrations (see code in Supplementary Section 13).

### DNA extraction, library preparation and sequencing

Samples of 2 mL were taken immediately after cold transport of the sludge, and centrifuged at 16,200 x *g* for 5 min at 4 °C to separate the biomass from the supernatant. The biomass pellets were stored at −80 °C until DNA extraction. The DNA of the 12 Nov 2020, 9 Jun, 16 Dec 2021 and 11 May 2022 samples was extracted with the DNeasy PowerSoil Pro Kit (Qiagen). The manufacturer’s instructions were followed, with exception of these steps: approximately 50 mg biomass was resuspended in the CD1 solution by vortexing before transferring to the PowerBead tube; 3 x 40 s bead-beating (Beadbeater-24, Biospec) was alternated with 2 min incubation on ice; tubes were gently inverted instead of vortexed to prevent DNA shearing ^35^. The DNA of the 20 Jan and 3 Mar 2021 samples (1/3 pellet) was extracted with the DNeasy UltraClean Microbial Kit (Qiagen) following the manufacturer’s instructions. The DNA concentration and quality were assessed with the Qubit 4 Fluorometer (Thermo Fisher Scientific) and the BioTek Synergy HTX multimode microplate reader (Agilent), respectively.

The samples of 12 Nov 2020 (np1), 9 Jun (np2), 16 Dec 2021 (np3) and 11 May 2022 (np4) were prepared for long-read sequencing using the Ligation Sequencing Kit V14 (Oxford Nanopore Technologies Ltd), the NEBNext^®^ Companion Module for Oxford Nanopore Technologies^®^ Ligation Sequencing (New England BioLabs) and UltraPure^TM^ BSA (50 mg/mL) (Thermo Fisher Scientific). The incubations in the Hula mixer were replaced with slow manual inversions, all resuspensions were performed by flicking the tube, and the last room temperature incubation step was performed a 37 °C to improve the recovery of long DNA fragments. Four MinION R10.4 flow cells (Oxford Nanopore Technologies), one for each sample, were used to sequence on a MinION for 89-96 h in accurate mode (260 bps), yielding 21-29 Gbp per sample. The sample of 20 Jan 2021 (np1.5) was prepared with a Ligation Sequencing Kit V12 and sequenced on a GridION with MinION R9.4 flow cells (Oxford Nanopore Technologies), generating 11.2 Gbp. Short-read sequencing was also performed on the samples of 20 Jan (il1) and 3 Mar 2021 (il2) on an Illumina NovaSeq 6000 platform by Novogene Ltd. (UK), resulting in over 20 Gbp (per sample) of 150 bp paired-end reads with a 350 bp insert.

### Processing of metagenomic data and MAG recovery

After sequencing, the DNA data was processed to obtain metagenome-assembled genomes (MAGs). The final set of MAGs was obtained from the five nanopore-sequenced samples (np1-4 and np1.5). The Illumina reads (il1 and il2) were solely used for differential coverage binning and to estimate the relative abundance of each MAG on the respective dates. The raw long reads were basecalled in super-accurate mode with the “dna_r10.4.1_450bps_sup.cfg” configuration file and -- do_read_splitting option using Guppy v6.4.2 (np1-4) or with “dna_r9.4.1_450bps_sup.cfg” using Guppy v5.0.7 (Oxford Nanopore Technologies) (np1.5). The duplex reads of np1-4 were filtered using pairs_from_summary and filter_pairs from Duplex tools v0.2.19 (Oxford Nanopore Technologies). The duplex reads were basecalled using the duplex basecaller of Guppy and merged with the remaining simplex reads using SeqKit v2.3.0 ^67^. The reads were filtered, trimmed and inspected with NanoFilt v2.8.0 ^68^ (options -q 10 −l 200), Porechop v0.2.4 (https://github.com/rrwick/Porechop) and NanoPlot v1.41.0 ^68^. The Illumina reads were filtered and trimmed using Trimmomatic v0.39 ^69^ with the options LEADING:3 TRAILING:3 SLIDINGWINDOW:4:15 MINLEN:35 HEADCROP:5. The kmer algorithm of Nonpareil v3.401 ^70^ estimated a diversity coverage of 69.9% (il1) and 71.3% (il2) for the trimmed Illumina reads.

The long reads were individually assembled and pairwise co-assembled (np1-np2, np2-np3, np3-np4) with Flye v2.9.1 ^71^ in --meta mode. The reads were mapped on the assembly with Minimap2 v2.24 ^72^. The individual assemblies were polished with Racon v1.4.3 (https://github.com/isovic/racon) and two times with Medaka v1.5.0 (https://github.com/nanoporetech/medaka). The reads from all samples were mapped to each assembly using Minimap2, the alignments were converted from SAM to BAM and sorted with SAMtools v1.10 ^73^ and the contig coverage in each sample was calculated with jgi_summarize_bam_contig_depths ^74^. The differential coverages were used for automatic binning of each assembly with MetaBAT2 v2.15 ^74^, MaxBin2 v2.2.7 ^75^ and CONCOCT v1.1.0 ^76^, setting the minimum contig length at 2000 bp. The outputs were combined into an optimized set of non-redundant bins with DAS Tool v1.1.3 ^77^, which used Prodigal v2.6.3 ^78^ and DIAMOND v2.0.8 ^79^. The bins obtained from all assemblies (np1, np1.5, np2, np3, np4, np1-np2, np2-np3, np3-np4) were dereplicated with the 1083 HQ MAGs from Singleton *et al.* (2021) ^35^ at 95% average nucleotide identity of open reading frames using dRep v3.2.2 ^80^ with the options -comp 70 -con 10 -sa 0.95 --S_algorithm gANI.

Bin completeness and contamination was assessed with the lineage_wf workflow of CheckM v1.1.3 ^81^. The relative abundance of the bins in each sample (np1, np2, np3, np4, il1, il2) was determined with CoverM v0.6.1 (https://github.com/wwood/CoverM), using the options -- methods relative_abundance mean --min-read-percent-identity 95 --min-read-aligned-percentage 50. Bins with completeness < 90%, contamination > 5% or with zero abundance in all samples were discarded, resulting in a non-redundant set of 349 HQ MAGs. The HQ MAGs were taxonomically classified using the classify_wf mode of GTDB-Tk v2.3.0 ^82^ and the GTDB release 207 ^83^ (<u>gtdbtk_r207_v2_data.tar.gz</u>). The presence of 16S rRNA genes was verified with barrnap v0.9 (https://github.com/tseemann/barrnap). A bacterial phylogenetic tree was made with FastTree v2.1.11 ^84^ using the multiple sequence alignment generated with the identify and align modes of GTDB-Tk, adjusted with the TreeTools v1.10.0 ^85^ package in RStudio v22.0.3 ^86^ with R v4.2.2 ^87^ and visualized with iTol v6.8.2 ^88^.

### Gene prediction and functional annotation

Genes were predicted in all assemblies using Prodigal v2.6.3 ^78^ with the -p meta option. The gene sequences were concatenated and duplicates were removed using grep and rmdup from SeqKit v2.3.1 ^67^, resulting in a unique set of genes covering all metagenomic samples. The predicted genes were functionally annotated with the annotate pipeline of EnrichM v0.6.5 (https://github.com/geronimp/enrichM), using DIAMOND v2.0.8 ^79^ and HMMER v3.2.1 (http://hmmer.org/) and the EnrichM v10 database, including a KO-annotated UniRef100 2018_11 ^89^ DIAMOND database and HMM libraries of the KEGG 88.2 ^90^, PFAM 32.0 ^91^, and TIGRFAMs 15.0 ^92^ databases. In general, the genes of interest from the nitrogen cycle were identified by their KO identifier (Table S3). Cytochrome P460 was identified through its PFAM identifier PF16694. The genes encoding the alpha- and beta-subunit of the cytoplasmic nitrate reductase (*narG* and *narH*) and the nitrite oxidoreductase (*nxrA* and *nxrB*) have the same KO identifier, so these were distinguished through a phylogenetic analysis using the graft command of GraftM ^93^ and the respective packages (7.70.nxrA_narG and 7.69.nxrB). If the alpha-subunit was classified as *narG* or *nxrA*, the putative beta-subunit located in the same contig was manually annotated. The unclassified sequences were left with the *narGH* annotation. The *nxrAB* genes from the *Ca.* Nitrotoga MAG (NOB) could not be distinguished with GraftM, but were confirmed with a BLAST on UniProt^94^. Similarly, the alpha-subunit of the ammonia monooxygenase gene (*amoA*) was distinguished from the methane monooxygenase gene (*pmoA*) using the 20170316_pmoA package of GraftM. Unidentified sequences remained annotated as *pmoA*. The beta- and gamma-subunits located in the same contig as *amoA* were manually annotated as *amoB* and *amoC*. Distinction between the quinol-dependent nitric oxide reductase (qNor, encoded by *norZ*) and the alpha subunit of the cytochrome c-dependent reductase (cNor, encoded by *norB*), was made by identifying the fused quinol oxidase domain on the N-terminal of *norZ* ^95^. A multiple sequence alignment was performed between putative NorB and NorZ protein sequences found in the metagenomes (K04561), and reference sequences of NorB (Pseudomonas stutzeri, P98008) and NorZ (Cupriavidus necator, Q0JYR9), extracted from UniProtKB ^94^, using Clustal Omega v1.2.4 ^96^. The alignment was visualized and analysed, and the quinol oxidase domain was identified with Jalview v2.11.3.2. The distinction between clade I and II nitrous oxide reductase, respectively TAT- and Sec-dependent, was made by combining the TIGRFAM annotation of EnrichM and the phylogenetic analysis of GraftM with the 7.45.nosZ package. The sequences not classified as either clade I or II remained annotated as unclassified *nosZ*. Data processing was performed using RStudio v22.0.3 ^86^ with R v4.2.2 ^87^, and the plyr v1.8.8 ^98^, tidyverse v2.0.0 ^99^, readxl v1.4.2 ^100^, data.table v1.15.0 ^101^, aplot v0.2.2 ^102^ and reshape2 v1.4.4 ^103^ packages.

### Protein extraction

Biomass samples were taken and stored as detailed in the DNA extraction section. Proteins were extracted from 12 samples, as previously described ^104^. Briefly, around 60 mg of the biomass pellet were homogenised in three cycles of vortexing and ice incubation with glass beads (150 – 212 µm, Sigma Aldrich), 50 mM TEAB buffer 1% (w/w) NaDOC and B-PER reagent (Thermo Scientific). Proteins in the supernatant were precipitated with 1:4 trichloroacetic acid (Sigma Aldrich). The pellet was washed and disrupted with acetone two times and re-dissolved in 200 mM ammonium bicarbonate with 6 M Urea (Sigma Aldrich). Human serum albumin (0.1 µg, Sigma Aldrich) was added to all samples to control the digestion efficiency. The mixture was reduced with 10 mM dithiothreitol (Sigma Aldrich) at 37 °C for 60 min, and alkylated with 20 mM iodoacetamide (Sigma Aldrich) in the dark for 30 min. Samples were diluted with 100 mM ammonium bicarbonate to obtain a urea concentration lower than 1 M. Protein digestion occurred overnight at 37 °C and 300 rpm with 1.5 µg sequencing grade trypsin (Promega). 0.5 pmol of the Pierce^TM^ Peptide Retention Time Calibration mix (Thermo Scientific) was added to all samples to control the chromatographic performance. Solid phase extraction was performed with an Oasis HLB 96-well µElution Plate (2 mg sorbent per well, 30 µm, Waters) and a vacuum pump. The columns were conditioned with MeOH, equilibrated with water two times, loaded with the peptide samples, washed with two rounds of 5% MeOH and sequentially eluted with 2% formic acid in 80% MeOH and 1 mM ammonium bicarbonate in 80% MeOH. The samples were dried in a centrifuge Concentrator plus (Eppendorf) at 45 °C and stored at −20 °C until analysis.

### Shotgun metaproteomics

Peptide samples were dissolved in 20 µL of 3% acetonitrile and 0.01% trifluoroacetic acid, incubated at room temperature for 30 min and vortexed thoroughly. The protein concentration was measured at 280 nm wavelength with a NanoDrop ND-1000 spectrophotometer (Thermo Scientific) and samples were diluted to a concentration of 0.5 mg/mL. Shotgun metaproteomics was performed as previously described ^104^, with a randomized sample order. Briefly, approximately 0.5 µg protein digest was analysed using a nano-liquid-chromatography system consisting of an EASY nano-LC 1200, equipped with an Acclaim PepMap RSLC RP C18 separation column (50 μm x 150 mm, 2 μm, Cat. No. 164568), and a QE plus Orbitrap mass spectrometer (Thermo Fisher Scientific). The flow rate was maintained at 350 nL/min over a linear gradient from 5% to 25% solvent B over 90 min, from 25% to 55% over 60 min, followed by back equilibration to starting conditions. Solvent A was a 0.1% formic acid solution in water (FA), and solvent B consisted of 80% ACN in water and 0.1% FA. The Orbitrap was operated in data dependent acquisition (DDA) mode acquiring peptide signals from 385–1250 m/z at 70 K resolution in full MS mode with a maximum ion injection time (IT) of 75 ms and an automatic gain control (AGC) target of 3E6. The top 10 precursors were selected for MS/MS analysis and subjected to fragmentation using higher-energy collisional dissociation (HCD) at a normalised collision energy of 28. MS/MS scans were acquired at 17.5 K resolution with AGC target of 2E5 and IT of 75 ms, 1.2 m/z isolation width. The protein reference sequence database was generated through whole metagenome sequencing of the microbial samples, which included all metagenome-assembled genomes (MAGs) and unique unbinned sequences from all samples. The raw mass spectrometric data from each sample were analysed against this database using PEAKS Studio X (Bioinformatics Solutions Inc.) in a two-round database search process. The initial round was conducted without considering variable modifications and missed cleavages. Subsequently, the focused database was further searched, allowing for a 20 ppm parent ion and a 0.02 m/z fragment ion mass error tolerance, up to 3 missed cleavages, and iodoacetamide as a fixed modification, with methionine oxidation and N/Q deamidation as variable modifications.

### Metaproteomic data analysis

Peptide spectrum matches were filtered against 5% false discovery rates (FDR) and protein identifications with ≥2 unique peptide sequences were considered significant. The human serum albumin added as internal process control was filtered out. Proteins were grouped according to their unique protein group identification. The peptide spectral counts were divided by their molar mass for normalisation and technical duplicates were averaged. The relative abundance of each protein in a certain sample was determined by dividing the respective normalized spectral counts by the sum of normalized spectral counts of all proteins detected in that sample. The total relative abundance of each MAG in the metaproteome was calculated by summing the relative abundance of all proteins belonging to that MAG. The same was performed to calculate the total relative abundance of functionally identical proteins. Some functionally identical proteins belonging to different MAGs from the same genus could not be distinguished because of their high similarity. Therefore, these proteins were grouped by their functional annotation and genus for the data analysis. Proteins that simultaneously matched unbinned sequences and one or more MAGs from a certain genus, were classified as belonging to that genus. The catalytic subunits of the nitrogen-converting enzymes of interest were used as representative of that protein during data analysis, with exception of the ammonia monooxygenase (AMO). The catalytic alpha-subunit (AmoA) is located in the cell membrane ^105^, and is thus hydrophobic, so it is not well detected in the proteomic analysis (Fig. S15). The beta-subunit (AmoB), only partially in the membrane, was detected in much higher amounts so it was here used as proxy for AMO. In any case, the results were similar for AmoA and AmoB (Fig. S17). Data processing was performed using RStudio v22.0.3 ^86^ with R v4.2.2 ^87^, and the plyr v1.8.8 ^98^, tidyverse v2.0.0 ^99^, readxl v1.4.2 ^100^, data.table v1.15.0 ^101^, aplot v0.2.2 ^102^, reshape2 v1.4.4 ^103^ and matrixStats v1.2.0 ^106^ packages.

## Supporting information

Full Supplementary Information

## Acknowledgements

We deeply thank Dirk Geerts (TU Delft) for valuable support with the bioreactor, Carol de Ram and Dita Heikens (TU Delft) for help with the protein extraction protocol, Waternet for providing the activated sludge (Hidde Schijfsma, Alex Veltman and Adrien Azé for sampling), Mads Albertsen and team at Aalborg University for sequencing one of the samples with Oxford Nanopore R9.4 during the “Hands-on metagenomics using Oxford Nanopore DNA sequencing” course, and Alexandra Deeke (Waterschap de Dommel), Cora Uijterlinde (STOWA), Inge Pistorius, Robert Kras, Floris de Heer and Maurice Ramaker (Waterschap Aa en Maas), Maaike Hoekstra (HHNK), Mariska Ronteltap (Hoogheemraadschap van Delfland), the Dutch Community of Practice for N_2_O, Adriano Joss (Eawag), and Wenzel Gruber (upwater AG) for insightful discussions.

## Data availability

Raw DNA reads were deposited in the NCBI Sequence Read Archive and the 54 high-quality MAGs were deposited in Genbank under BioProject PRJNA1082082. The raw mass spectrometry proteomics data acquired in this project have been deposited in the ProteomeXchange Consortium database under the dataset identifier PXD051095. The HQ MAGs accession numbers, quality and abundance can be found in Supplementary Data 1, and the gene presence and protein abundance in the MAGs and the unbinned sequences can be found in Supplementary Data 2 and 3, respectively. The Python codes used to calculate the maximum N_2_O rates and to simulate the biological nitrogen removal process are in Supplementary Information.

## Author contributions

NR, MvL and ML conceptualized the study. NR, MP, MvD, CHM, MvL and ML designed the experiments. NR, MvD and CHM performed the maximum activity measurements. MZ collected and pre-processed the WWTP data. NR performed the Nanopore sequencing, implemented the bioinformatics pipeline and performed metagenomic analysis with input from TA. NR performed the protein extraction with input from MP. MP performed the shotgun metaproteomics. NR wrote the Python and RStudio scripts for data analysis and NR, MvD, CHM analysed the data with inputs from MZ, MvL and ML. NR wrote the draft manuscript and created the visuals with strong inputs from ML and contributions from all co-authors. All authors reviewed and approved the final manuscript.

