## Supplementary material for "Long-term multi-meta-omics resolves the ecophysiological controls of seasonal N_2_O emissions": Full Supplementary Information

### 1. Summary of literature on seasonal $N_2O$ emissions in WWTPs

**Table S1. Summary of literature on worldwide seasonal $N_2O$ emission events in WWTPs.** Information of the WWTPs and the monitoring campaigns: location, type of WWTP (SBR = sequencing batch reactor; AI = intermittent aeration; BNR = biological nitrogen removal; OD = oxidation ditch; CarR = carrousel reactor; Den = denitrification; Nit = nitrification; PF = plug-flow; GS = granular sludge; A2O = anaerobic-anoxic-oxic; AO = anoxic-oxic; MBR = membrane bioreactor; CAS = conventional activated sludge), $N_2O$ monitoring period, water temperature (T), dissolved $O_2$ (DO), and the period of seasonal $N_2O$ emissions. The main conclusions of each paper – observations, hypotheses and main $N_2O$ producing pathways (NN = nitrifier nitrification/hydroxylamine oxidation; ND = nitrifier denitrification; HD = heterotrophic denitrification) – are also described. All plants were activated sludge (AS) plants except Dinxperlo WWTP (granular sludge).

| Country | WWTP | Monitoring period | T (ºC) | DO (mg/L) | Seasonal $N_2O$ emissions | Observations/Hypotheses | $N_2O$ pathway | Ref. |
|---|---|---|---|---|---|---|---|---|
| **Southern hemisphere (Winter: Jun-Sep)** | | | | | | | | |
| Australia (Adelaide) | SBR A/I | $N_2O$: Feb – Mar 2014<br>$NO_2^-$: Jan 2014 – Jun 2017 | - | 0.5-2.8 | No long-term $N_2O$ data | 3x $NO_2^-$ peak in Jun – Sep.<br>No long-term $N_2O$ measurement, so no hypotheses for seasonal peak. | - | 1 |
| Brasil (Rio de Janeiro) | Non-BNR | Jan – Jul (< 1 year) | 24-32 | 0-7 | Jan - Apr | Simultaneous high $NO_2^-$. Strong positive correlation $N_2O$ vs. T. Hypothesis: AOB outcompete NOB at higher T, leading to $NO_2^-$ accumulation and $N_2O$ emissions. | - | 2 |
| **Northern hemisphere (Winter: Dec-Mar)** | | | | | | | | |
| China (Beijing) | OD (a), A2O (b), reversed A2O (c) | Mar-Nov (< 1 year) | - | - | Mar - Jun (a), Mar-Apr (b), Jun-Jul (c) | No hypotheses for $N_2O$ seasonality. | - | 3 |
| Denmark (Avedøre WWTP) | CarR AI | Mar 2018 – Feb 2019 | 10-20 | 0.5-1.5 | 3x Mar - July | $N_2O$ peaks during increasing T, but likely not main cause. Negative correlation $N_2O$ vs. DNA abundance of $N_2O$ reducers and NOB (16S rRNA genes). | NN | 4,5 |
| Finland (Viikinmäki WWTP) | Den/Nit | Jul 2012 – Jun 2013 | 9-21 | 3.5 [a] | ↑ Emissions in winter and spring | No clear relationship between T and $N_2O$ emissions. | - | 6 |
| Finland | Den/Nit | Jan 2019 – Nov 2022 | 10-18 | - | Dec-Apr | $NO_2^-$ linked to winter $N_2O$ emissions. Hypothesis: seasonal emissions partly due to T. | NN/ND | 7 |
| Netherlands (Kralingseveer WWTP) | PF (Den/Nit) + carR (Den/Nit) | Oct 2010 – Jan 2012 | 10-20 | 0.5-2 | Feb - Jun | Positive correlation $N_2O$ vs. maximum $NO_2^-$ concentration (negatively correlated with T). Seasonal $N_2O$ lags 2-3 months behind T. $N_2O$ correlated with $NH_4^+$, $NO_2^-$ and $NO_3^-$. | ND/HD | 8–10 |
| Netherlands (Dinxperlo WWTP) | GS Nereda® | Aug 2017 – Mar 2018 (< 1 year) | 8-22 | 0-3 | Dec – Feb | $NO_2^-$ in effluent remained low. | ND/HD | 11 |
| South Korea (3 WWTPs in Gwangju) | A2O (a), AO MBR (b), SBR (c) | Apr 2018 – Jan 2019 (< 1 year) | 15-30 | 4.3 ± 2.2 | Not clear, ↑ emissions in Aug (a), Apr (b), Dec (c) | DIC/VSS and $sOUR_{AOB}$ (*ex situ*) positively correlated with $N_2O$ in the aerobic AS. DOC/$NO_x^-$ negatively correlated with $N_2O$ in the anoxic AS. | NN/HD | 12 |
| South Korea (Gockseong) | SBR (only Nit) | Apr 2018 – Jul 2019 | 9-30 | > 3 | Dec - Mar | Seasonal $NO_2^-$ accumulation. ↑ $sOUR_{AOB}/sOUR_{NOB}$ (*ex situ*) during $N_2O$ peak. Hypothesis: difference between AOB and NOB activity results in $NO_2^-$ and $N_2O$ emissions. | - | 13 |
| Switzerland (3 WWTPs) | AI (a), CAS (b), SBR (c) | Mar 2014 – Sep 2015 (a)<br>Dec 2015 – Mar 2017 (b)<br>Feb 2018 – Apr 2019 (c) | 10-20 (a), 12-23 (b), 12-21 (c) | 2 (a,b), 2-3 (c) | Dec-Mar (a), Dec – Apr (b), Jan – May (c) | Simultaneous $NO_2^-$ accumulation. ↓ NOB and filamentous bacteria (16S rRNA genes) during peaks. Hypothesis: compromised floc integrity led to NOB washout and consequent $NO_2^-$ and $N_2O$ accumulation. Increasing SRT and DO did not improve the plant performance. [b] | - | 14,15 |
| Switzerland (5 WWTPs) | AO (a,b), A2O (c), SBR (d), AI (e) | > 1 year | - | - | Jan-Apr (a), Apr-Jul (c), No seasonality (b,d,e) | WWTPs with seasonal $N_2O$ had higher emission factor. Hypothesis: reduced NOB performance causes $NO_2^-$ and $N_2O$ accumulation. Propose all-year denitrification (no nitrification-only periods) to avoid $NO_2^-$ accumulation. | ND/HD | 16 |

[a] Average values
[b] Measures were only applied as contingency instead of prevention, it may have been too late.

### 2. Full-scale WWTP configuration and operational parameters

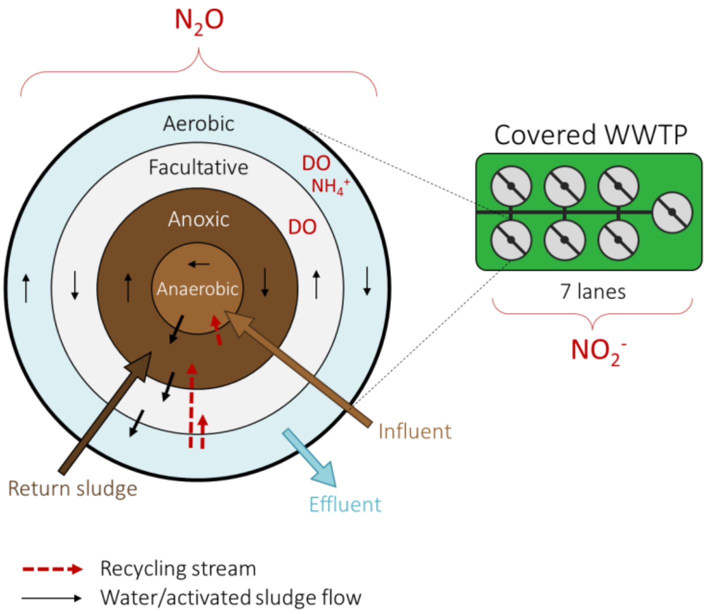

**Figure S1. Schematic representation of the configuration of the monitored activated sludge reactor.** The dissolved oxygen (DO) and $NH_4^+$ concentrations were measured in the aerobic compartment, the $N_2O$ was measured in the combined off-gas of all compartments, and $NO_2^-$ was measured in the pooled effluent of all seven lanes of the WWTP.

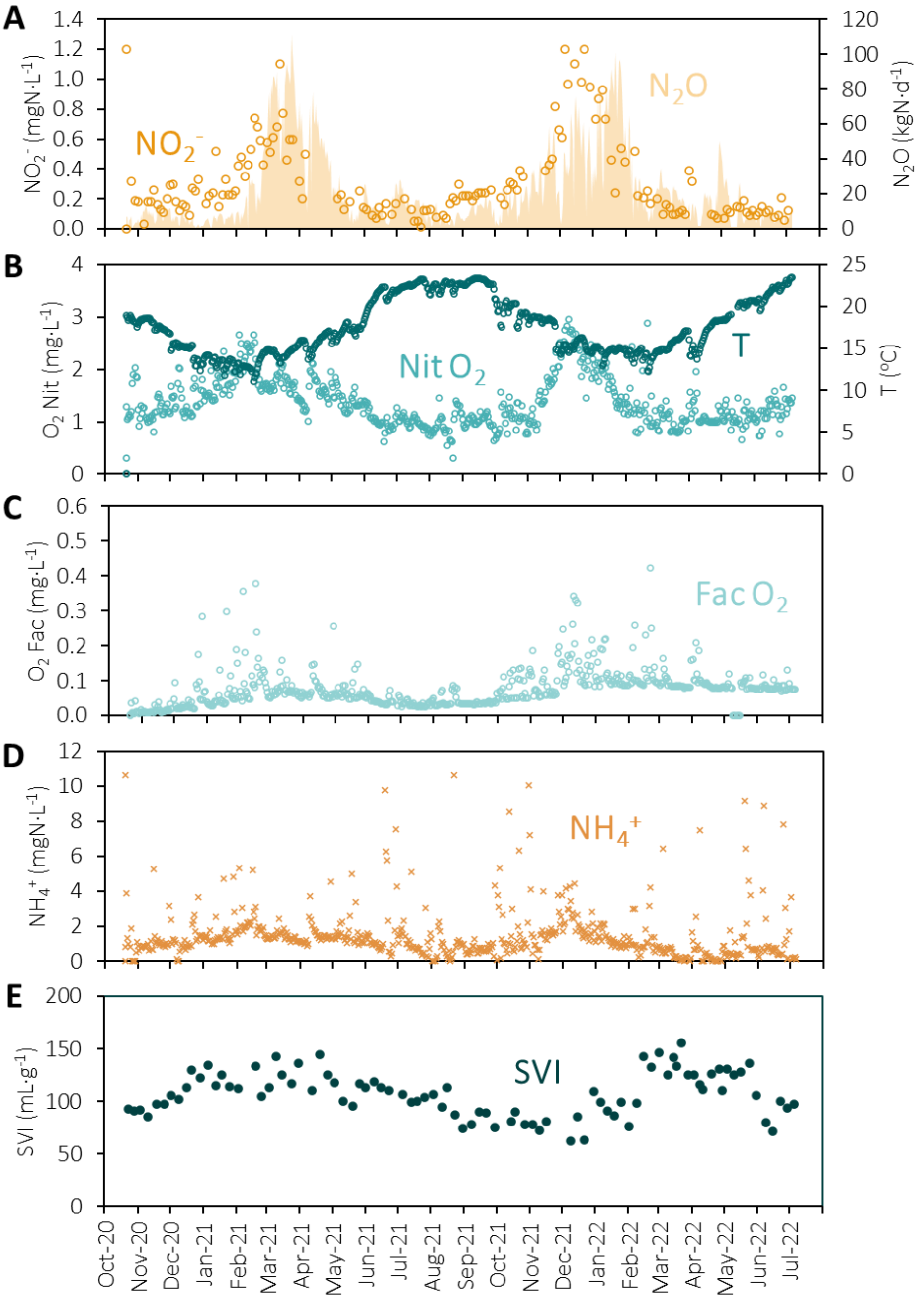

**Figure S2. Daily averages of the wastewater treatment plant (WWTP ) parameters between Oct 2020 and Jul 2022. (A)** Nitrite concentration in the effluent (seven lanes pooled together, symbols, left axis) and $N_2O$ emission rates measured from off-gas measurements of all compartments of a single covered biological nutrient removal lane of the WWTP (area, right axis). **(B)** Dissolved $O_2$ concentration in the aerated compartment (light blue, left axis) and water temperature inside the reactor (dark blue, right axis). **(C)** Dissolved $O_2$ concentration in the facultative compartment, which depends on the nitrogen removal performance of the system. **(D)** $NH_4^+$ concentration in the nitrification compartment. Outliers above 12 mg $N \cdot L^{-1}$ were omitted for clarity. High $NH_4^+$ concentrations often coincided with rainy days. All metabolites were measured in a single biological nutrient removal lane of the WWTP, except the effluent $NO_2^-$ (seven lanes pooled together). **(E)** Sludge volume index representing the settleability of the sludge.

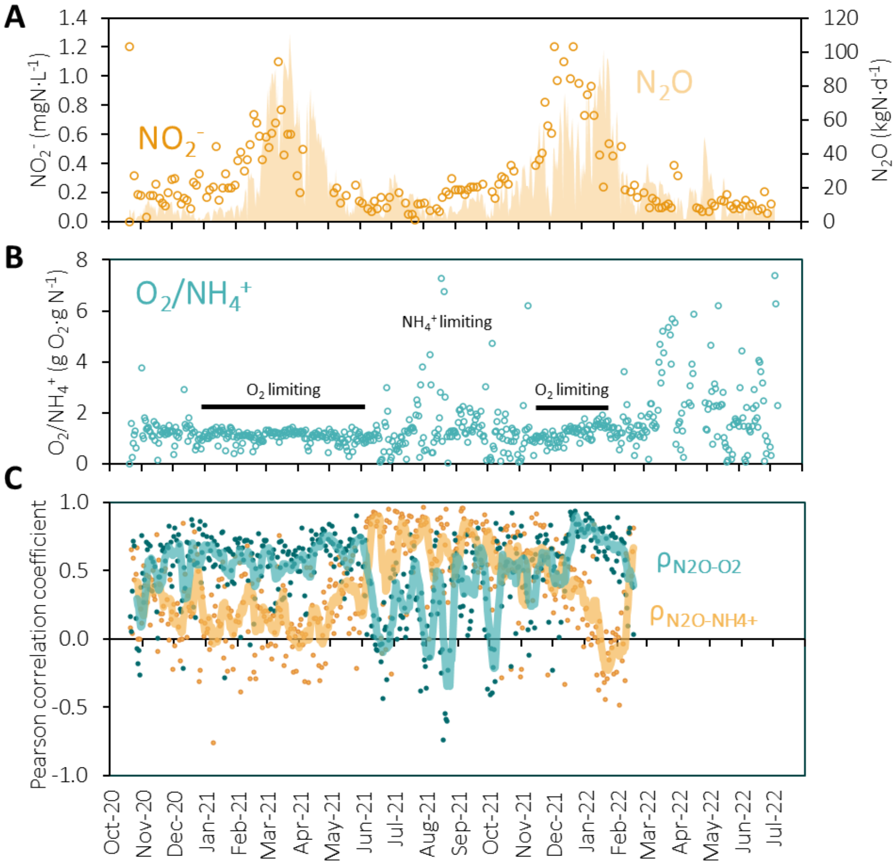

**Figure S3. Seasonal variations in $O_2$ and $NH_4^+$ limiting conditions. (A)** Daily averages: Nitrite concentration in the effluent (seven lanes pooled together, symbols, left axis) and $N_2O$ emission rates measured from off-gas measurements of all compartments of a single covered biological nutrient removal lane of the WWTP (area, right axis). **(B)** Ratio between the daily average dissolved $O_2$ and $NH_4^+$ concentrations in the nitrification compartment. During the seasonal nitrogen oxides peak the $O_2/NH_4^+$ reaches a plateau, suggesting that the $O_2$ is limiting. In the summer months this plateau is not observed, suggesting that there is enough $O_2$ to fully consume the influent $NH_4^+$. Outliers above 8 g $O_2$·g $N^{-1}$ were omitted for clarity. **(C)** The Pearson correlation coefficient (ρ) between the continuously measured (every 15 min) concentrations of $N_2O$-$O_2$ (blue symbols) and $N_2O$-$NH_4^+$ (yellow symbols) was calculated for each day. The weekly averages are represented with a line. The high values for $N_2O$-$O_2$ in winter suggest a direct dependence of $N_2O$ production on the $O_2$ concentration, supporting $O_2$ as the rate-determining (limiting) substrate during the nitrogen oxides peaks. The opposite is observed in summer, with mainly the $NH_4^+$ concentrations determining the $N_2O$ production.

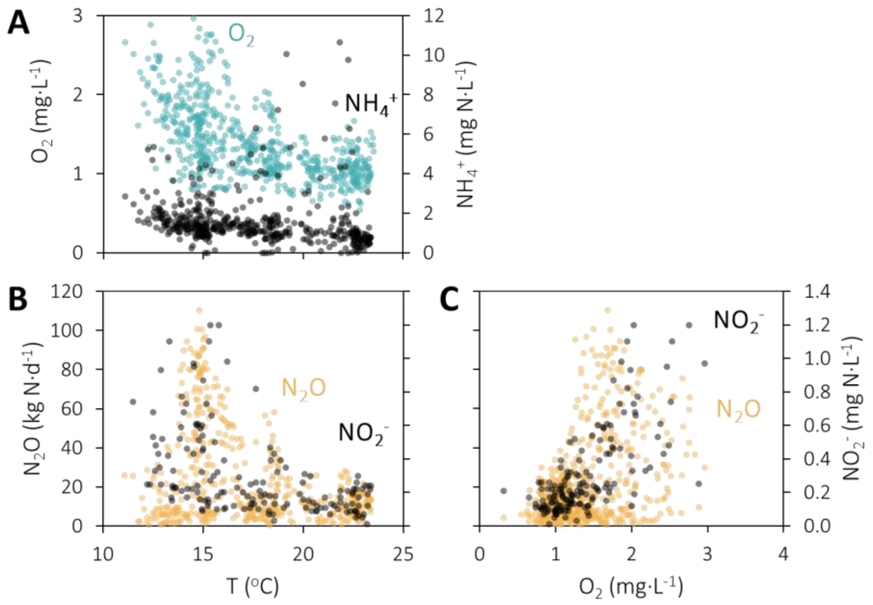

**Figure S4. Correlation between key parameters of the full-scale WWTP. (A)** Dissolved $O_2$ concentration (blue, left axis) and $NH_4^+$ concentration in the nitrification compartment (black, right axis) as function of the water temperature. **(B)** $N_2O$ emission rate (orange, left axis) and effluent $NO_2^-$ concentration (black, right axis) as function of the water temperature. **(C)** $N_2O$ emission rate (orange, left axis) and effluent $NO_2^-$ concentration (black, right axis) as function of the dissolved $O_2$ concentration in the nitrification compartment.

**Table S2. Pearson correlation coefficients between $NO_2^-$ and $N_2O$ and the weekly averages of several WWTP parameters:** $N_2O$ emission rate, nitrite concentration in the effluent, temperature in the nitrification tank, total suspended solids (TSS), ammonium concentration in the nitrification tank, nitrate concentration in the nitrification tank, dissolved oxygen concentration in the nitrification tank and the sludge volume index. Negative correlations are highlighted in red, and positive correlations are highlighted in green. Strong correlations are highlighted in bold.

| | $NO_2^-$ | T | TSS | $NH_4^+$ | $NO_3^-$ | $O_2$ | SVI |
|---|---|---|---|---|---|---|---|
| $N_2O$ | **0.7** | -0.4 | -0.2 | 0.1 | 0.0 | 0.5 | 0.0 |
| $NO_2^-$ | | -0.5 | 0.0 | 0.4 | 0.0 | **0.8** | -0.2 |

### 3. Maximum nitrifying and denitrifying activities

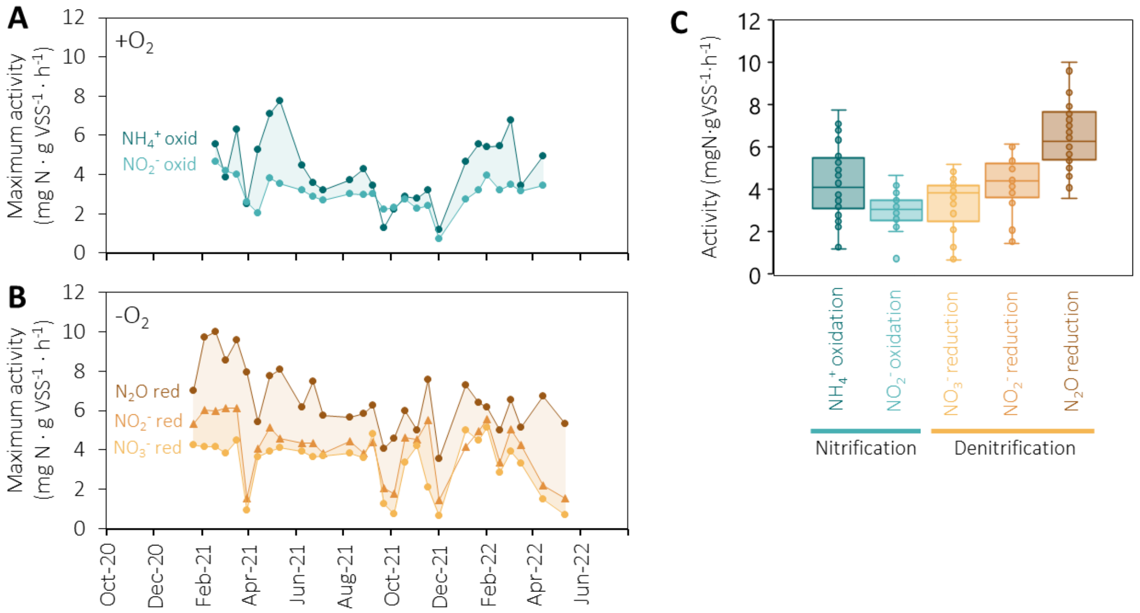

**Figure S5.** ***Ex situ*** **maximum nitrifying and denitrifying activity of activated sludge measured at 20 ºC. (A)** $NH_4^+$ and $NO_2^-$ oxidation measured under oxic conditions. **(B)** $N_2O$, $NO_2^-$ and $NO_3^-$ reduction measured under anoxic conditions. **(C)** Boxplots summarizing the *ex situ* nitrifying and denitrifying activities of activated sludge.

### 4. Metagenomic data processing

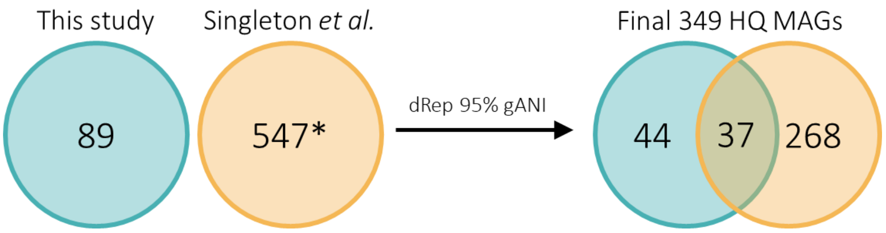

**Figure S6. Number of HQ MAGs before and after dereplication with the MAGs from Singleton *et al.* [17] at 95% average nucleotide identity of open reading frames (gANI).** In the final set of HQ MAGs, there were 37 overlapping MAGs between our original set and the original Singleton *et al.* set, from which 10 of our MAGs and 27 of Singleton *et al.*'s MAGs were kept by the dereplication software. So, in total, the final set contained 54 of our MAGs and 295 of Singleton *et al.*'s MAGs. *After dereplicating the Singleton *et al.* 1083 MAGs at 95% gANI and before filtering out MAGs that were not present in our samples (the 252 MAGs from Singleton *et al.* with zero abundance in our samples were filtered out after dereplication).

**Table S3. Nitrogen metabolism genes and KO identifiers.**

| KO ID | Gene | Description |
|---|---|---|
| K10944 | amoA | methane/ammonia monooxygenase subunit A [EC:1.14.18.3 1.14.99.39] |
| K10945 | amoB | methane/ammonia monooxygenase subunit B |
| K10946 | amoC | methane/ammonia monooxygenase subunit C |
| K10535 | hao | hydroxylamine dehydrogenase [EC:1.7.2.6] |
| K00370 | narG, narZ, nxrA | nitrate reductase / nitrite oxidoreductase, alpha subunit [EC:1.7.5.1 1.7.99.-] |
| K00371 | narH, narY, nxrB | nitrate reductase / nitrite oxidoreductase, beta subunit [EC:1.7.5.1 1.7.99.-] |
| K00374 | narI, narV | nitrate reductase gamma subunit [EC:1.7.5.1 1.7.99.-] |
| K02567 | napA | nitrate reductase (cytochrome) [EC:1.9.6.1] |
| K02568 | napB | nitrate reductase (cytochrome), electron transfer subunit |
| K00368 | nirK | nitrite reductase (NO-forming) [EC:1.7.2.1] |
| K15864 | nirS | nitrite reductase (NO-forming) / hydroxylamine reductase [EC:1.7.2.1 1.7.99.1] |
| K04561 | norB | nitric oxide reductase subunit B [EC:1.7.2.5] |
| K02305 | norC | nitric oxide reductase subunit C |
| K00376 | nosZ | nitrous-oxide reductase [EC:1.7.2.4] |
| K03385 | nrfA | nitrite reductase (cytochrome c-552) [EC:1.7.2.2] |
| K15876 | nrfH | cytochrome c nitrite reductase small subunit |
| K20932 | hzsA | hydrazine synthase alpha subunit [EC:1.7.2.7] |
| K20933 | hzsB | hydrazine synthase beta subunit [EC:1.7.2.7] |
| K20934 | hzsC | hydrazine synthase gamma subunit [EC:1.7.2.7] |
| K20935 | hdh | hydrazine dehydrogenase [EC:1.7.2.8] |

### 5. Temporal profiles of the MAGs abundance in terms of DNA and protein

The data presented in this section can be found in Supplementary Data 1.

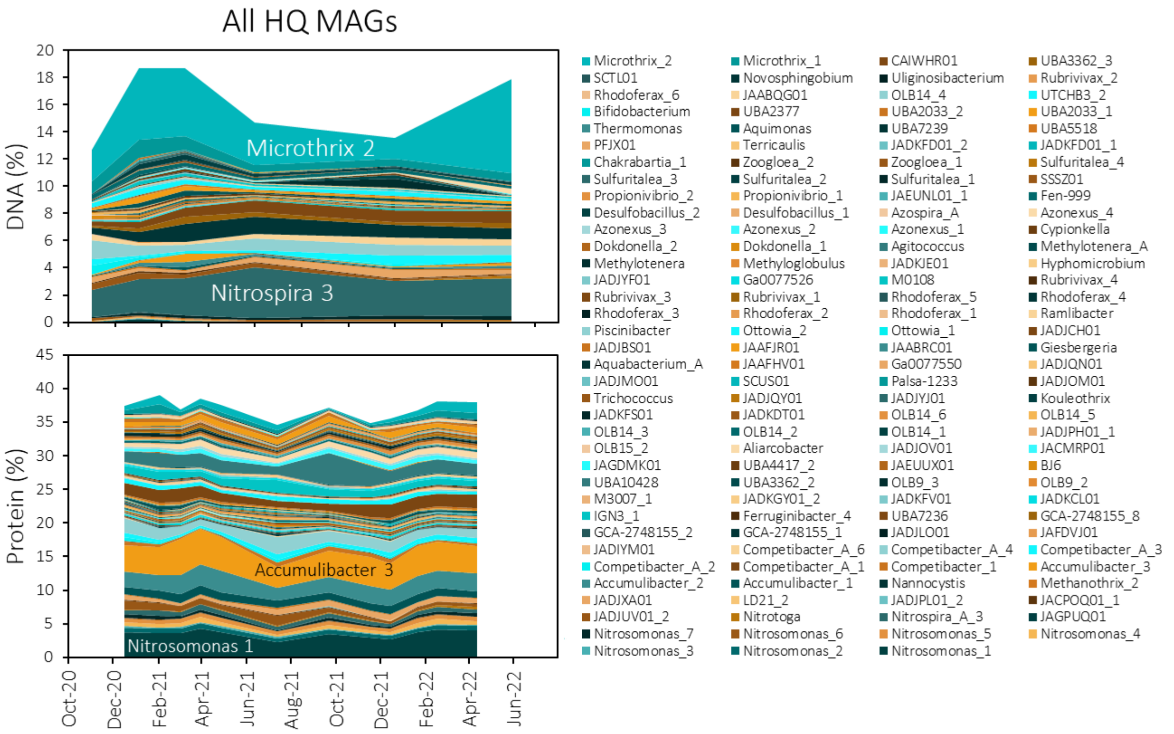

**Figure S7. Temporal fluctuations in the relative abundance of the 143 HQ MAGs detected in the proteome in terms of DNA and protein.**

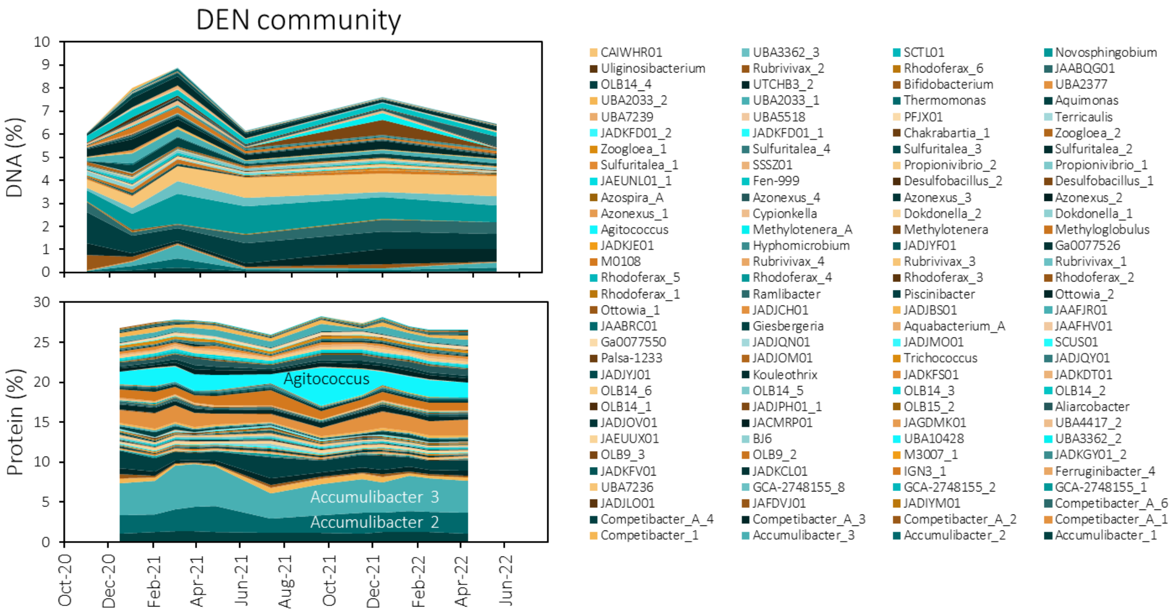

**Figure S8. Temporal fluctuations in the relative abundance of the 124 DEN MAGs detected in the proteome in terms of DNA and protein.**

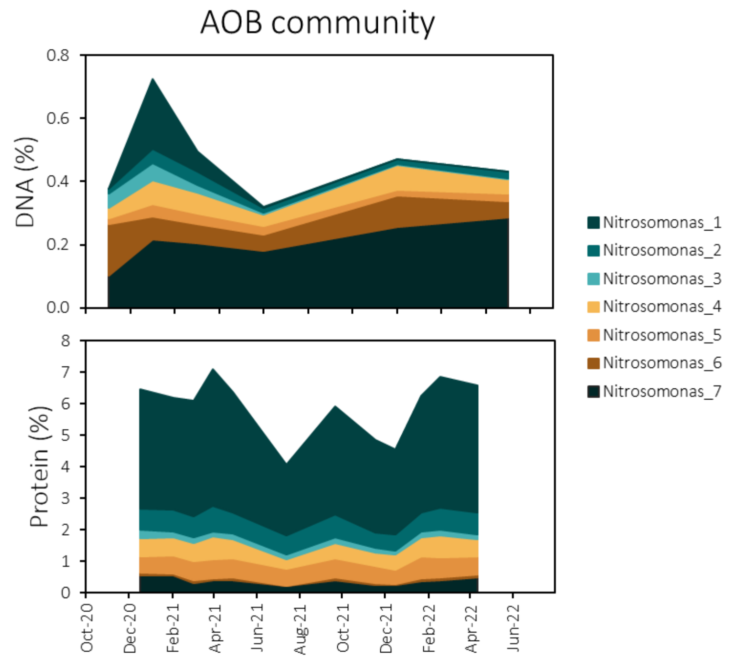

**Figure S9. Temporal fluctuations in the relative abundance of the seven AOB MAGs in terms of DNA and protein.**

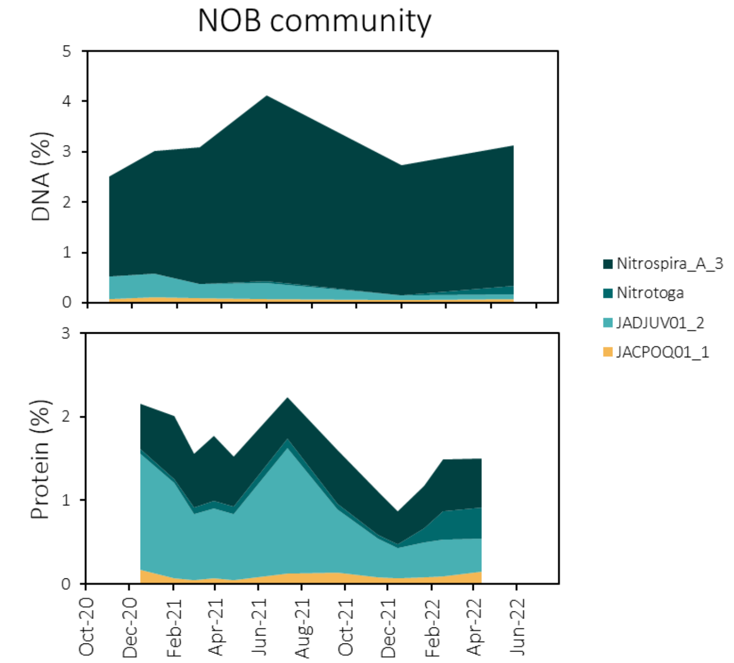

**Figure S10. Temporal fluctuations in the relative abundance of the four NOB MAGs detected in the proteome in terms of DNA and protein.**

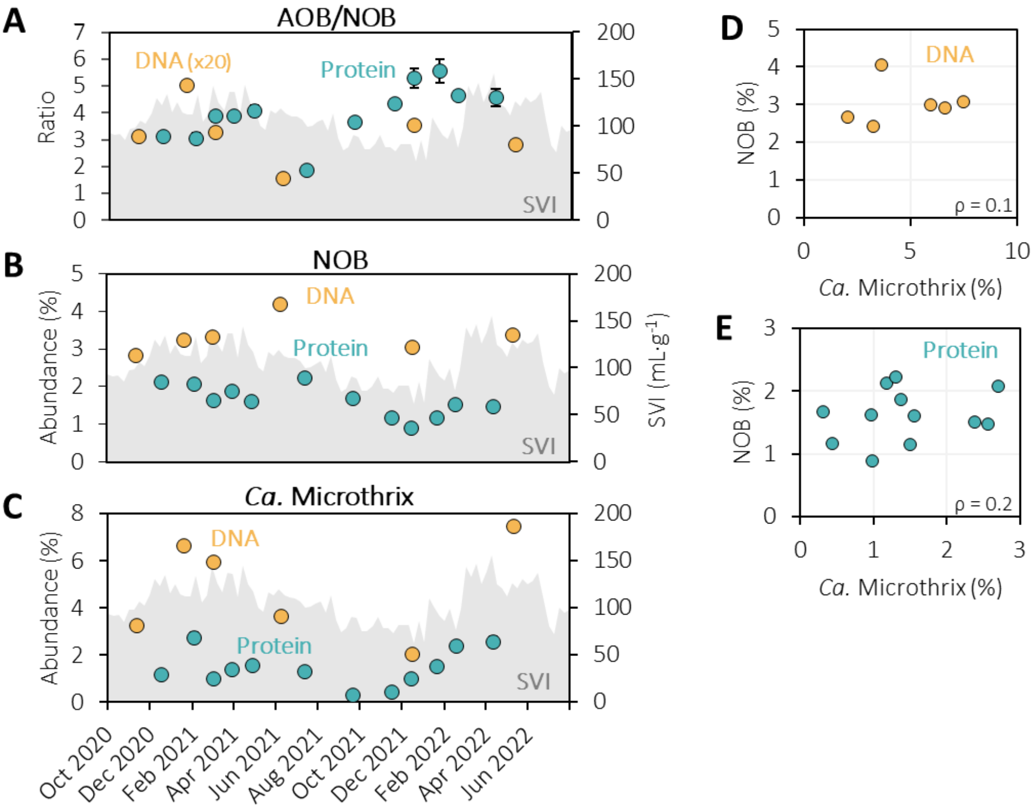

**Figure S11. Relative DNA and protein abundances of nitrifying and filamentous bacteria alongside the sludge volume index. Left axis: (A)** Ratio AOB/NOB; **(B)** NOB abundance; **(C)** *Ca.* Microthrix (filamentous) abundance. The error bars in all protein ratios and abundances were propagated from standard deviations of technical duplicates and most are smaller than the symbols. Right axis: **(A-C)** Sludge volume index (SVI), representing the sludge settleability. Higher SVI stands for a worse settleability. High amounts of filamentous bacteria cause bad settleability (high SVI), this is also seen in this plot. **(D, E)** There was no correlation between the abundances of NOB and *Ca.* Microthrix (ρ = Pearson correlation coefficient).

**Table S4. Pearson correlation coefficients between the protein abundance of all genera with an abundance above 0.1% and weekly average WWTP operational parameters.** Twelve timepoints were used to calculate the correlation coefficients. The genera are ordered from high to low abundance. Correlation coefficients below -0.7 or above 0.7 are highlighted.

| | | Pearson correlation coefficient | | | | | |
|---|---|---|---|---|---|---|---|
| **Type** | **Genus** | **T** | **$NH_4^+$** | **DO** | **$N_2O$** | **$NO_2^-$** | **SVI** |
| DEN | Accumulibacter | -0.5 | 0.0 | 0.2 | 0.6 | 0.0 | 0.5 |
| AOB | Nitrosomonas | -0.6 | -0.3 | -0.2 | 0.0 | -0.4 | 0.7 |
| DEN | Competibacter | 0.4 | 0.0 | -0.1 | -0.5 | 0.0 | -0.5 |
| DEN | Agitococcus | 0.5 | -0.3 | -0.2 | -0.1 | 0.0 | -0.3 |
| DEN | JADJCH01 | -0.8 | 0.4 | 0.6 | 0.4 | 0.5 | 0.1 |
| DEN | Azonexus | -0.1 | -0.2 | 0.0 | 0.6 | 0.1 | 0.1 |
| Other | Microthrix | -0.6 | 0.5 | 0.1 | -0.3 | -0.3 | 0.6 |
| DEN | UBA2033 | 0.2 | 0.0 | 0.1 | -0.1 | 0.1 | 0.0 |
| DEN | M0108 | 0.5 | 0.1 | -0.3 | -0.3 | -0.2 | -0.2 |
| DEN | Sulfuritalea | -0.3 | -0.1 | 0.0 | 0.1 | -0.1 | 0.4 |
| NOB | JADJUV01 | 0.2 | -0.1 | -0.3 | -0.5 | -0.3 | 0.1 |
| NOB | Nitrospira | -0.1 | -0.2 | -0.2 | -0.2 | -0.4 | 0.6 |
| Other | JADJXA01 | 0.6 | -0.2 | -0.2 | -0.2 | 0.2 | -0.6 |
| DEN | OLB14 | 0.5 | -0.1 | -0.3 | -0.4 | -0.2 | -0.2 |
| DEN | Ottowia | 0.1 | 0.1 | 0.3 | -0.1 | 0.5 | -0.5 |
| DEN | Rhodoferax | -0.7 | 0.4 | 0.4 | 0.2 | 0.3 | 0.4 |
| DEN | Propionivibrio | -0.1 | 0.2 | 0.2 | 0.3 | 0.1 | 0.1 |
| DEN | Zoogloea | -0.3 | 0.5 | 0.7 | 0.6 | 0.5 | 0.0 |
| DEN | Giesbergeria | -0.8 | 0.5 | 0.7 | 0.4 | 0.5 | 0.0 |
| DEN | SSSZ01 | 0.1 | 0.1 | 0.2 | 0.1 | 0.2 | -0.2 |
| DEN | JAEUUX01 | 0.4 | 0.2 | 0.3 | 0.3 | 0.6 | -0.4 |
| DEN | Rubrivivax | -0.9 | 0.2 | 0.5 | 0.2 | 0.3 | 0.3 |
| DEN | Ga0077526 | -0.4 | 0.6 | 0.5 | 0.2 | 0.1 | 0.2 |
| DEN | JAABQG01 | -0.7 | 0.3 | 0.2 | -0.4 | 0.0 | 0.4 |
| DEN | Fen-999 | -0.5 | 0.1 | 0.5 | 0.6 | 0.4 | 0.0 |
| DEN | Ramlibacter | -0.6 | 0.2 | 0.0 | -0.1 | -0.4 | 0.6 |
| DEN | PFJX01 | 0.2 | -0.3 | -0.1 | 0.2 | 0.1 | -0.2 |
| DEN | JAEUNL01 | 0.1 | 0.4 | 0.4 | 0.5 | 0.3 | -0.4 |
| DEN | JADJBS01 | -0.6 | 0.6 | 0.6 | -0.1 | 0.6 | 0.0 |
| DEN | Hyphomicrobium | 0.1 | 0.2 | -0.2 | -0.2 | -0.2 | 0.2 |
| DEN | JADJYF01 | 0.0 | 0.6 | 0.6 | 0.0 | 0.5 | -0.1 |
| DEN | Dokdonella | -0.3 | 0.3 | 0.4 | 0.0 | 0.5 | -0.4 |
| DEN | JADKCL01 | 0.2 | 0.1 | 0.1 | 0.4 | 0.3 | 0.0 |
| DEN | JAAFJR01 | -0.6 | -0.2 | -0.2 | 0.1 | -0.2 | 0.6 |
| DEN | JADJVA01 | 0.2 | -0.1 | 0.1 | 0.3 | 0.4 | 0.0 |
| NOB | Nitrotoga | -0.2 | -0.2 | -0.4 | -0.2 | -0.4 | 0.4 |
| DEN | JADKGY01 | -0.2 | -0.2 | 0.1 | 0.4 | 0.1 | 0.3 |
| DEN | Palsa-1233 | 0.0 | 0.3 | 0.4 | 0.5 | 0.4 | 0.2 |
| DEN | GCA-2748155 | 0.2 | 0.5 | 0.1 | -0.4 | -0.1 | -0.1 |
| DEN | Piscinibacter | -0.6 | 0.3 | 0.2 | 0.2 | 0.3 | 0.4 |
| DEN | OLB9 | -0.1 | 0.3 | -0.1 | 0.0 | 0.0 | 0.4 |
| DEN | IGN3 | 0.0 | -0.1 | 0.1 | 0.2 | 0.2 | 0.2 |
| DEN | UBA5518 | -0.4 | 0.2 | 0.1 | -0.4 | -0.1 | 0.3 |

### 6. Taxa quantification in terms of DNA and protein

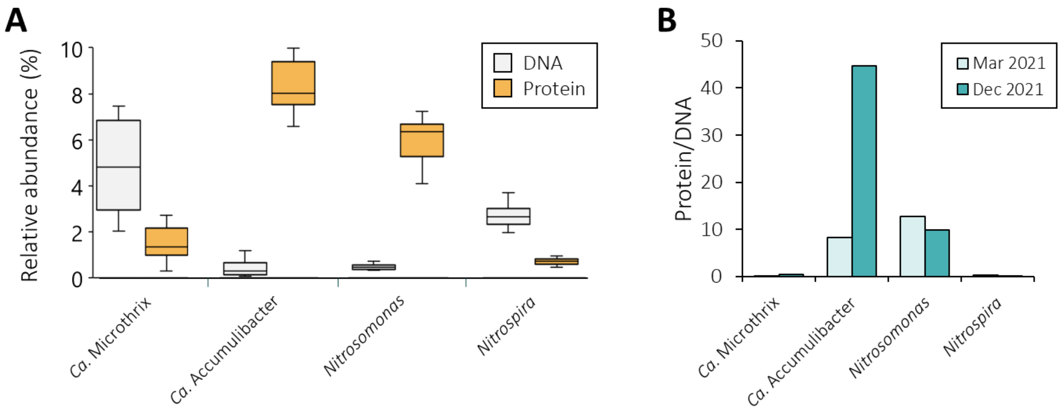

**Figure S12. DNA and proteomic quantification of key activated sludge genera. (A)** Overview of the relative abundance of *Ca.* Microthrix (filamentous bacteria), *Ca.* Accumulibacter (phosphate-accumulating organisms), *Nitrosomonas* (AOB) and *Nitrospira* (NOB), as quantified in the DNA (6 samples) and protein (12 samples). **(B)** Protein over DNA ratio of the same genera, measured in metagenomic and metaproteomic samples taken on the same day.

A disparity between genomic and proteomic quantification was observed for several key genera: *Ca.* Microthrix (filamentous), *Ca.* Accumulibacter (phosphate-accumulating), *Nitrosomonas* (AOB) and *Nitrospira* (NOB). Specifically, as often observed in wastewater treatment systems [5,13,19,21,22], the relative contribution of AOB *Nitrosomonas* to the community's DNA pool was 7-fold lower than NOB. The opposite is expected since AOB have a higher biomass yield per g nitrogen than NOB (Table S8). The under- or over-representation of certain taxa in the sludge DNA pool is likely an artifact of the quantification of cell number (DNA) vs. active biomass (protein) [18–20]. The relatively higher protein-based quantification of AOB compared to NOB likely reflects the larger AOB cell size (Table S5, Fig. S13).

**Table S5. The cell volumes of several AOB and NOB species were estimated based on cell sizes described in literature.**

| Guild | Species | D (µm) | L (µm) | Estimated volume (µm$^3$)* | Reference |
|---|---|---|---|---|---|
| AOB | *Nitrosomonas europaea* | 0.8-1.1 | 1.0-1.7 | 0.50-1.6 | 23 |
| | *Nitrosomonas eutropha* | 1.0-1.3 | 1.6-2.3 | 1.3-3.1 | 23 |
| | *Nitrosomonas halophila* | 1.1-1.5 | 1.5-2.2 | 1.4-3.9 | 23 |
| | *Nitrosomonas mobilis* | 1.5-1.7 | 1.5-2.1 | 2.7-4.8 | 23 |
| | *Nitrosomonas communis* | 1.0-1.4 | 1.7-2.2 | 1.3-3.4 | 23 |
| NOB | *Nitrospira marina* | 0.3-0.4 | 0.8-1.0 | 0.06-0.13 | 24 |
| | *Nitrospira moscoviensis* | 0.2-0.4 | 0.9-2.2 | 0.03-0.28 | 25 |
| | *Nitrospira defluvii* | 0.2-0.4 | 0.7-1.7 | 0.02-0.21 | 26 |
| | *Nitrospira calida* | 0.3-0.5 | 1.0-2.2 | 0.07-0.43 | 27 |
| | *Ca.* Nitrospira inopinata | 0.2-0.3 | 0.7-1.6 | 0.02-0.11 | 28 |

*Assuming cylindrical shape: $V = \pi \cdot L \cdot D^2/4$

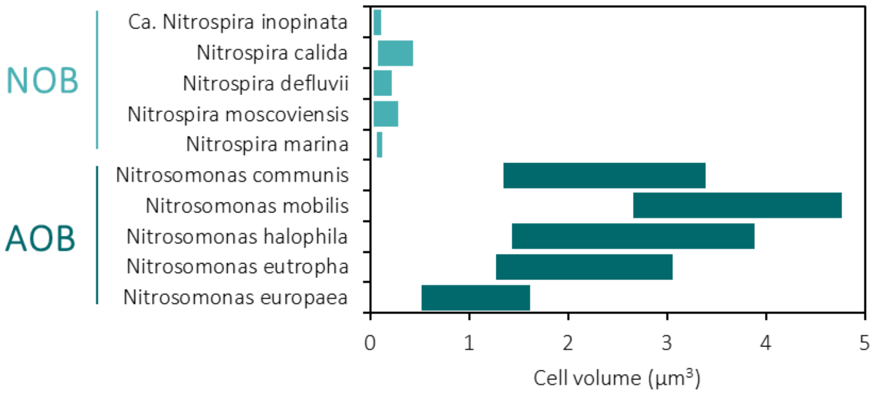

**Figure S13. Estimated cell volumes of *Nitrospira* (light blue) and *Nitrosomonas* species (dark blue).**

### 7. Functional characterization of the HQ MAGs

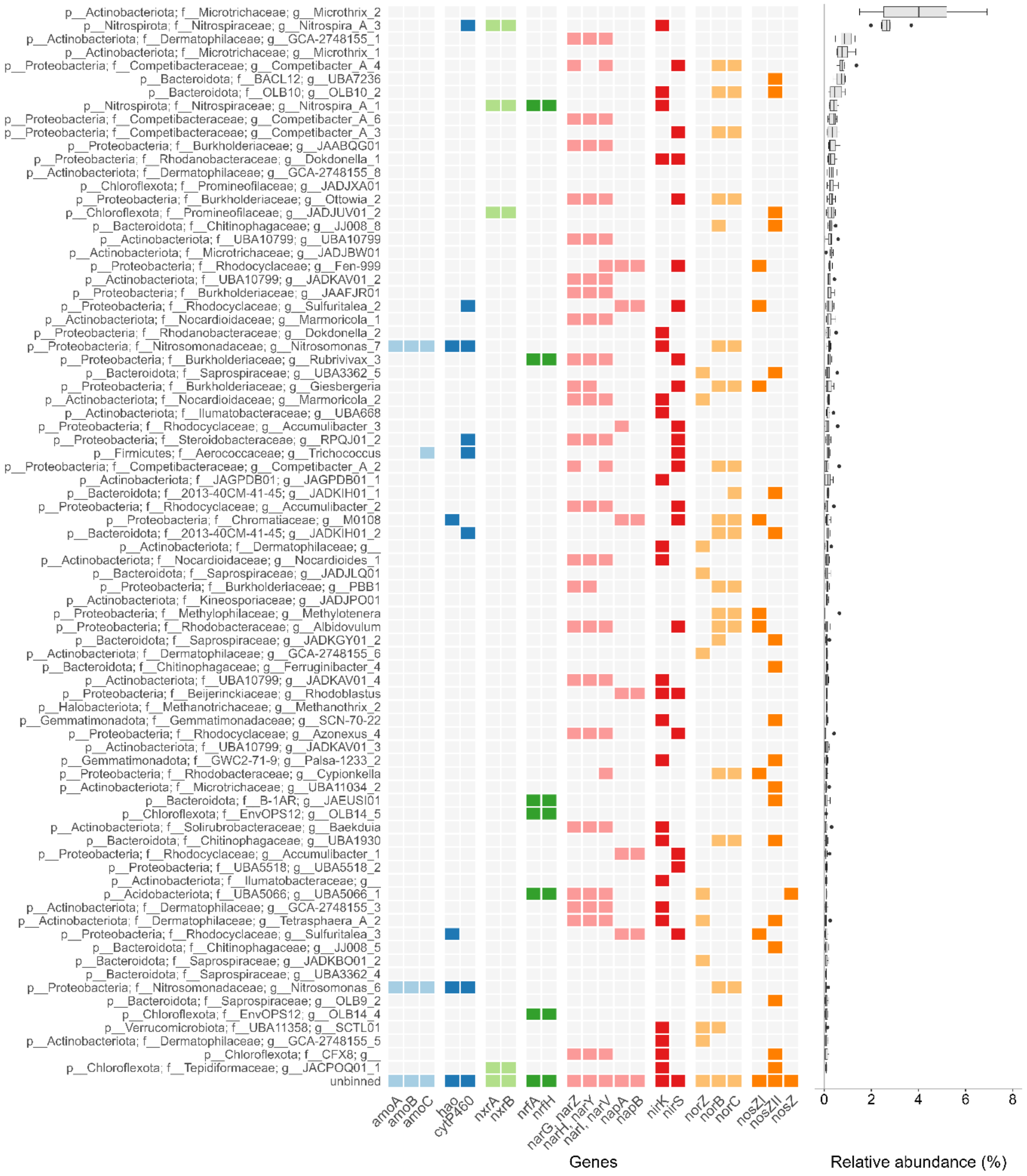

**Figure S14. Nitrogen gene content of the HQ MAGs with an average relative abundance above or equal to 0.8%.** The MAGs are ordered from most to least abundant and are identified by the phylum, family and genus. The heatmap shows the presence (colored) or absence (light grey) of nitrogen cycle genes: ammonia oxidation (light blue), hydroxylamine oxidation (dark blue), nitrite oxidation (light green), dissimilatory nitrite reduction to ammonia (dark green), nitrate reduction (pink), nitrite reduction (red), nitric oxide reduction (light orange) and nitrous oxide reduction (dark orange). The *nosZ* genes were identified as clade I (*nosZI*), clade II (*nosZII*) or unknown (*nosZ*). The boxplot on the right represents the relative abundance of each MAG at the six time points that were sequenced. The abundance of the unbinned fraction (72%) was omitted for clarity. The full set of nitrogen cycle genes in the HQ MAGs is in Supplementary Data 2.

### 8. Protein expression profiles

The relative abundance of the nitrogen enzymes in all twelve proteomic samples can be found in Supplementary Data 2.

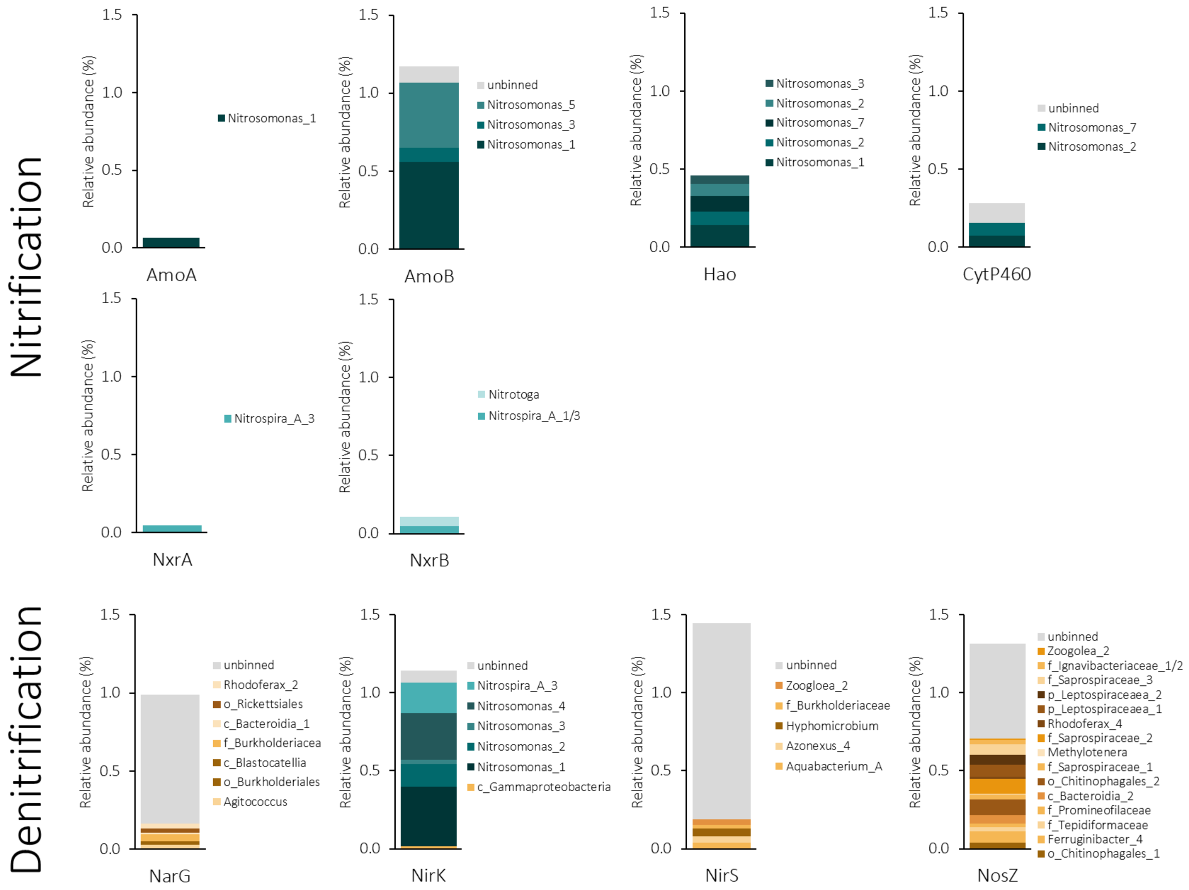

**Figure S15. Average protein abundances (12 samples) and distribution per MAG.** The top two rows represent the enzymes involved in the oxidation of ammonium to nitrite and nitrate: alpha- and beta-subunits of the ammonia monooxygenase (AmoA and AmoB), hydroxylamine oxidoreductase (Hao), the hydroxylamine oxidising cytochrome P460 (CytP460) and the alpha- and beta- subunits of the nitrite oxidoreductase (NxrA and NxrB). The bottom row represents the enzymes involved in the reduction of nitrogen oxides to dinitrogen gas: catalytic subunit of the nitrate reductase (NarG), copper- and cd1-type nitrite reductase (NirK and NirS) and catalytic subunit of the nitrous oxide reductase (NosZ). The coloured portion of the bars correspond to proteins belonging to a certain MAG (blue to nitrifiers and yellow to denitrifiers) and the grey part corresponds to proteins belonging to the unbinned portion of the community. **Note:** enzymes are divided based on the reaction they perform, the guilds are distinguished by colour. For example, NirK is effectively a denitrifying enzyme but it was almost entirely expressed by nitrifiers (blue).

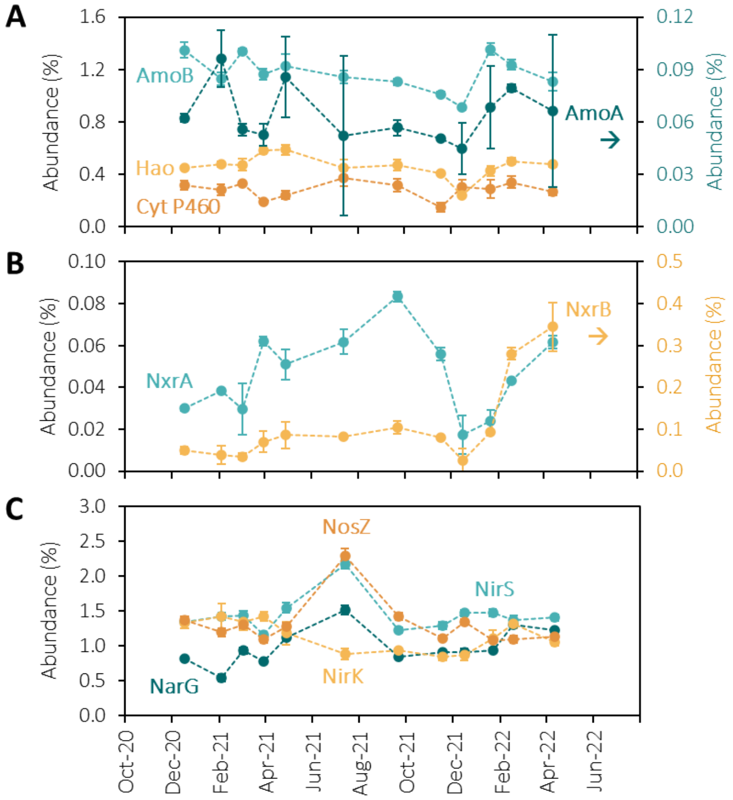

**Figure S16. Relative abundance of enzymes involved in ammonia (A) and nitrite oxidation (B), and in the reduction of nitrate, nitrite and nitrous oxide (C). (A)** Ammonia oxidation to nitrite: Alpha and beta subunits of the ammonia monooxygenase (AmoA on the right axis, and AmoB on the left axis), hydroxylamine oxidoreductase (Hao, left axis), and the cytochrome P460 (Cyt P460, left axis). **(B)** Nitrite oxidation to nitrate: Alpha and beta subunits of the nitrite oxidoreductase (NxrA on the left axis and NxrB on the right axis). **(C)** Nitrate reduction to dinitrogen gas: catalytic subunit of the membrane-bound nitrate reductase (NarG), Cu-type nitrite reductase (NirK), *cd1*-type nitrite reductase (NirS) and nitrous oxide reductase (NosZ). All abundances include MAG and unbinned proteins. The error bars in all protein ratios were propagated from standard deviations of technical duplicates.

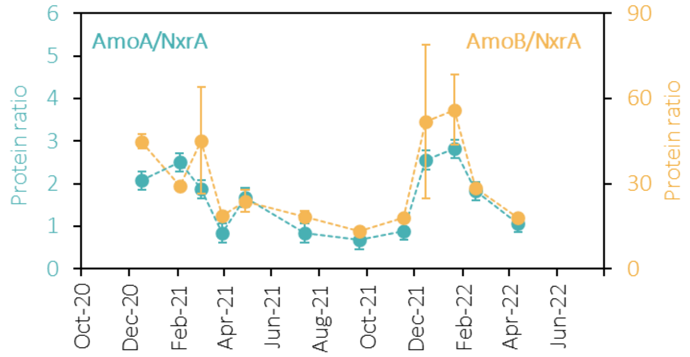

**Figure S17. Identical protein ratio profiles when using either the alpha or the beta subunits of AMO.** The beta subunit of the ammonia monooxygenase (AmoB) was used in all our analyses because the catalytic alpha subunit (AmoA) was detected in very low amounts. In any case, the protein profiles using either of the subunits is identical. All abundances include MAG and unbinned proteins. The error bars in the protein ratios were propagated from standard deviations of technical duplicates.

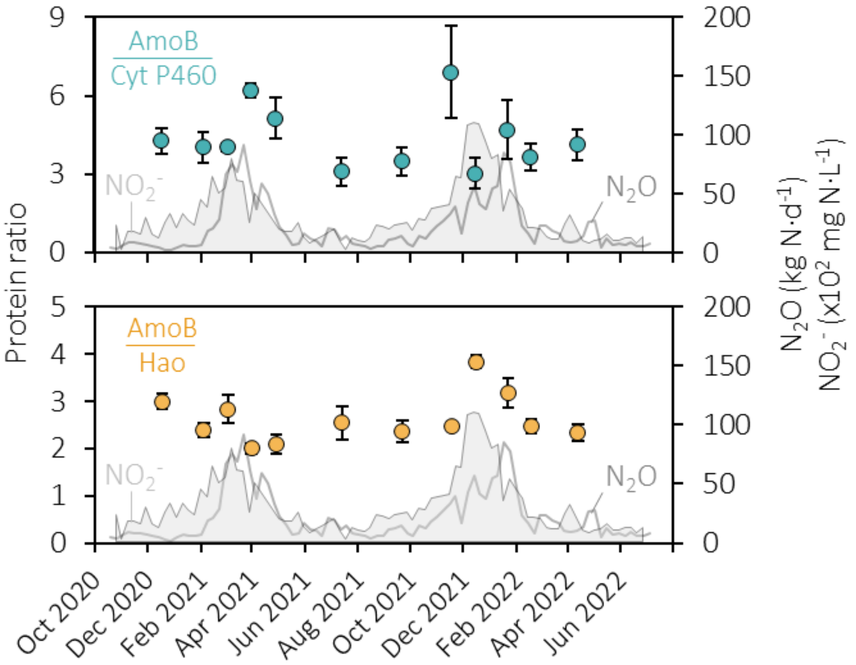

**Figure S18. Fluctuation of the ratio between hydroxylamine-producing (AmoB) and -consuming (Cyt P460 and Hao) enzymes.** The ratio between the beta-subunit of the ammonia monooxygenase (AmoB) and cytochrome P460 (Cyt P460) and the hydroxylamine oxidoreductase (Hao) represent the hydroxylamine flux balance. All abundances include MAG and unbinned proteins. The error bars in all protein ratios were propagated from standard deviations of technical duplicates.

### 9. Balance of DEN at proteomic and kinetic levels

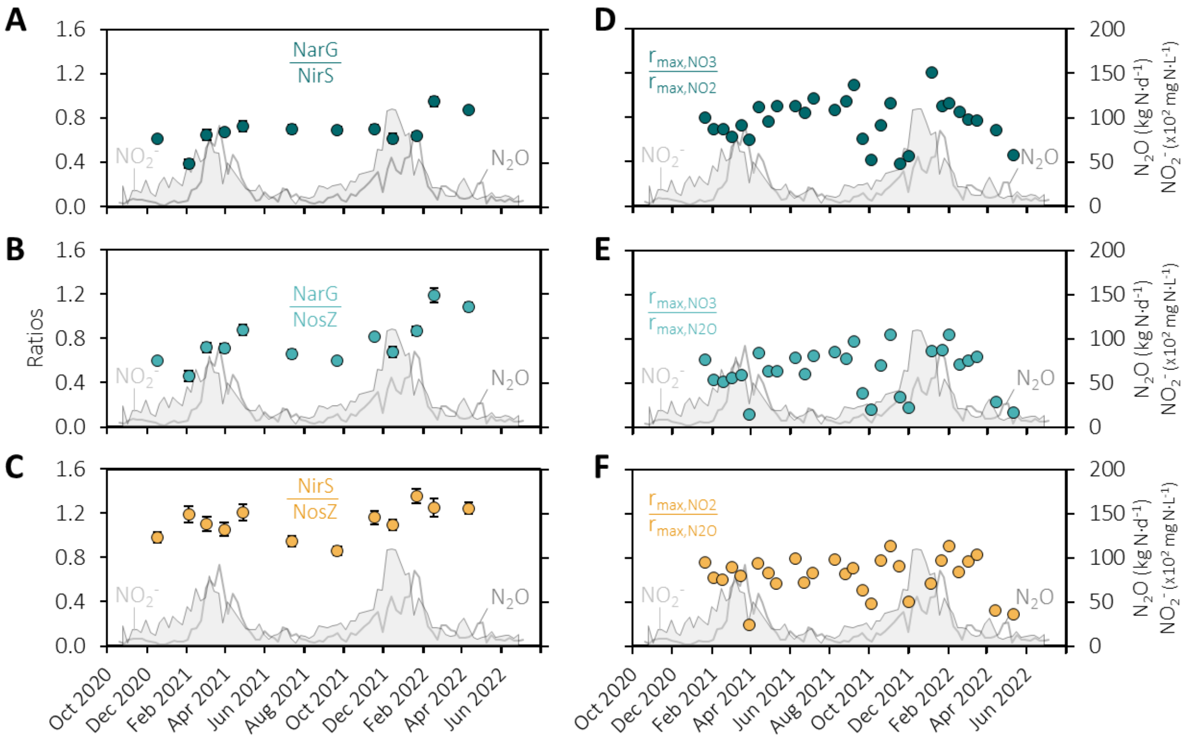

**Figure S19. Fluctuations in the balance of DEN fluxes in terms of protein abundances (A-C) and maximum activities (D-F). (A)** Ratio between the nitrite-producing and -consuming enzymes of DEN: membrane-bound nitrate reductase (NarG) and *cd1*-type nitrite reductase (NirS). **(B)** Ratio between membrane-bound nitrate reductase (NarG) and nitrous oxide reductase (NosZ). **(C)** Ratio between the $N_2O$-producing and -consuming enzymes of DEN: NirS and NosZ. The error bars in all protein ratios were propagated from standard deviations of technical duplicates and some are smaller than the symbols. All abundances include MAG and unbinned proteins. **(D)** Ratio between the nitrate and nitrite reduction potentials. **(E)** Ratio between the nitrate and nitrous oxide reducing potentials. **(F)** Ratio between the nitrite and nitrous oxide reduction potentials.

### 10. Free ammonia and free nitrous acid toxicity

The maximum possible concentration of free ammonia (FA, $NH_3$) and free nitrous acid (FNA, $HNO_2$) in the WWTP was calculated using the maximum pH and $NH_4^+$ concentration for FA and the minimum pH and maximum $NO_2^-$ concentration for FNA.

$$C_{NH_3} = 10^{pH-pKa} \cdot C_{NH_4^+} \quad \text{(eq. S1)}$$

$$C_{HNO_2} = 10^{pKa-pH} \cdot C_{NO_2^-} \quad \text{(eq. S2)}$$

**Table S6. Values used to calculate the maximum free ammonia concentration in the WWTP and the literature reported thresholds for AOB and NOB.**

| Maximum pH | pKa | Maximum $NH_4^+$ (mg N·L$^{-1}$) | Maximum $NH_3$ (mg N·L$^{-1}$) | Threshold AOB [29] (mg $NH_3$-N·L$^{-1}$) | Threshold NOB [29] (mg $NH_3$-N·L$^{-1}$) |
|---|---|---|---|---|---|
| 6.7 | 9.26 | 11 | 0.03 | 10 | 0.1 |

**Table S7. Values used to calculate the maximum free nitrous acid concentration in the WWTP and the literature reported thresholds for AOB and NOB.**

| Minimum pH | pKa | Maximum $NO_2^-$ (mg N·L$^{-1}$) | Maximum $HNO_2$ (mg N·L$^{-1}$) | Threshold NOB [30] (mg $HNO_2$-N·L$^{-1}$) | Threshold NOB [30] (mg $HNO_2$-N·L$^{-1}$) |
|---|---|---|---|---|---|
| 6.15 | 3.16 | 1.2 | 0.001 | 0.2 | 0.01 |

### 11. AOB and NOB growth stoichiometry and kinetics

**Table S8. Maximum specific growth rates and biomass yields of *Nitrosomonas* and *Nitrospira* cultures reported in literature.** The $\mu_{max}$ values were normalized to 20 °C using the Arrhenius equation (eq. S5) and the experimentally determined coefficients for AOB and NOB (reported at the end of this page).

| Guild | Species | $\mu_{max}$ ($d^{-1}$) | $Y_{X/N}$ (gX/gN) | Conditions | $\mu_{max}$ 20 °C ($d^{-1}$) | Reference |
|---|---|---|---|---|---|---|
| AOB | *Nitrosomonas europaea* | 0.84 | 0.063 | 30 °C, pH 7 | 0.27 | 31 |
| | *Nitrosomonas spp.* | 1.0 | 0.12 | 30 °C, pH 7 | 0.32 | 32 |
| | *Nitrosomonas spp.* | 0.54 | 0.14 | 24 °C, pH 7.8 | 0.34 | 33 |
| | *Nitrosomonas europaea* | 1.3 | - | 30 °C, pH 6.8 | 0.42 | 34 |
| | *Nitrosomonas europaea* | 1.3 | 0.36 | 28 °C, pH 7.9 | 0.53 | 35 |
| Median | | 1.00 | 0.13 | | 0.34 | |
| NOB | *Nitrospira defluvii* | 0.64 | 0.017* | 28 °C, pH 7.5 | 0.30 | 36 |
| | *Nitrospira moscoviensis* | 0.75 | 0.030* | 37 °C, pH 7.5 | 0.15 | 36 |
| | *Nitrospira sp.* | 0.64 | 0.020* | 28 °C, pH 7.5 | 0.30 | 36 |
| | *Nitrospira sp.* | 0.32 | - | 29 °C, pH 8.0 | 0.14 | 37 |
| | *Nitrospira japonica* | 0.62 | - | 29 °C, pH 8.0 | 0.26 | 37 |
| | *Nitrospira spp.* | 0.69 | 0.09 | 22 °C, pH 7.5 | 0.57 | 38 |
| Median | | 0.64 | 0.025 | | 0.28 | |

*Assuming a protein content of 50% the dry mass

The growth rates of AOB and NOB depend on the temperature and substrate concentration:

$$\mu_{AOB} = \mu_{max,AOB}(T) \cdot \frac{c_{NH_4^+}}{c_{NH_4^+}+K_{NH_4^+}} \cdot \frac{c_{O_2}}{c_{O_2}+K_{O_2,AOB}} \quad \text{(eq. S3)}$$

$$\mu_{NOB} = \mu_{max,NOB}(T) \cdot \frac{c_{NO_2^-}}{c_{NO_2^-}+K_{NO_2^-}} \cdot \frac{c_{O_2}}{c_{O_2}+K_{O_2,NOB}} \quad \text{(eq. S4)}$$

With $\mu$ the specific growth rate, $\mu_{max}$ the maximum specific growth rate (a function of temperature), $c_i$ the concentrations of $NH_4^+$, $NO_2^-$ and $O_2$ and $K_i$ the half-saturation constants for each substrate.
The maximum specific growth rates depend on the temperature according to the Arrhenius equation:

$$\mu_{max,i} = \mu_{max,i}^{T_{ref}} \cdot \theta_i^{(T-T_{ref})}, \quad i = AOB, NOB \quad \text{(eq. S5)}$$

With $\theta$ the Arrhenius coefficient and $T_{ref}$ a reference temperature for which we know $\mu_{max}$.

The Arrhenius coefficient for AOB and NOB was determined from five independent experiments with activated sludge, sampled between October 2021 and February 2022. The maximum $NH_4^+$ and $NO_2^-$ oxidation rates were determined at 10, 15, 20 and 25 °C. The Arrhenius coefficients, obtained from linear regressions of eq. S5 (Fig. S20), were 1.12 ± 0.01 for AOB and 1.10 ± 0.01 for NOB.

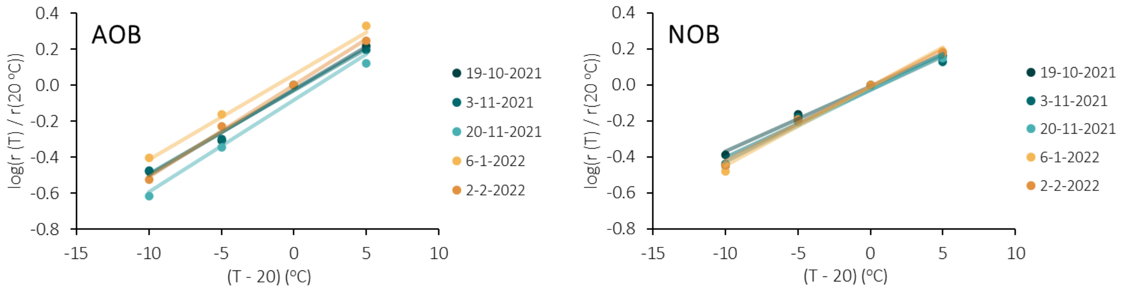

**Figure S20. Linear regressions between maximum activities and temperature to determine the Arrhenius coefficients of AOB and NOB in activated sludge.**

The $\mu_{max}$ of AOB and NOB in the activated sludge was estimated from the maximum $NH_4^+$ and $NO_2^-$ oxidation activities ($r_i$ in $gN\cdot L^{-1}\cdot d^{-1}$), the median biomass yield on substrate retrieved from literature ($Y_{X/N}$ in $gX\cdot gN^{-1}$, Table S8), the median relative abundance of AOB and NOB in our samples in terms of protein ($p_i$ in %), and the median total biomass concentration ($c_X$ as volatile suspended solids, $gVSS\cdot L^{-1}$).

$$\mu_{max,i} = \frac{r_i \cdot Y^i_{X/N}}{p_i \cdot c_X}, \quad i = AOB, NOB \qquad \text{(eq. S6)}$$

The obtained values of $\mu_{max}$ were 0.20 (AOB) and 0.12 $d^{-1}$ (NOB). These values are comparable to the ones reported in literature (Table S8).

In the WWTP, when the growth rates of AOB and NOB drop below 1/SRT the bacteria cannot grow as fast as they are removed from the system, so they start to washout, i.e. their concentration in the sludge slowly decreases. The variation of $\mu_{max}$ with temperature was determined with eq. S5 and represented in Fig. S21. The temperature at which each guild starts to wash out, here named critical temperature (Tc), is higher for NOB than AOB (Fig. S21). This means that during winter, when the temperature progressively decreases, NOB starts washing out before AOB, resulting in a higher AOB/NOB ratio.

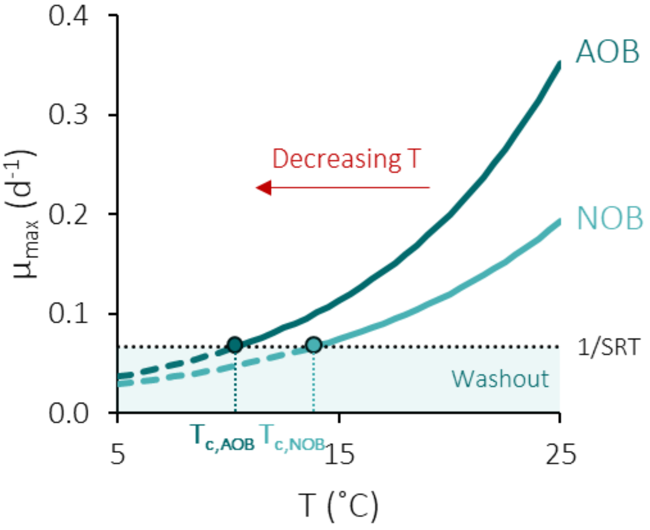

**Figure S21. Variation of the $\mu_{max}$ of AOB and NOB with a decreasing temperature from 25 to 5 ˚C.** The critical temperature ($T_c$) represents the temperature below which each guild starts washing out.

### 12. Mathematical model replicating the seasonal peaks

A simplified mathematical model describing the microbial and metabolite dynamics in a biological nutrient removal process was set up in Python v3.9.12 (Fig. S22). The goal was to replicate the seasonal nitrifier and nitrogen metabolites profiles. The metabolic cascade was replicated simply by changing the temperature and the oxygen concentration, as occurs in the WWTP.

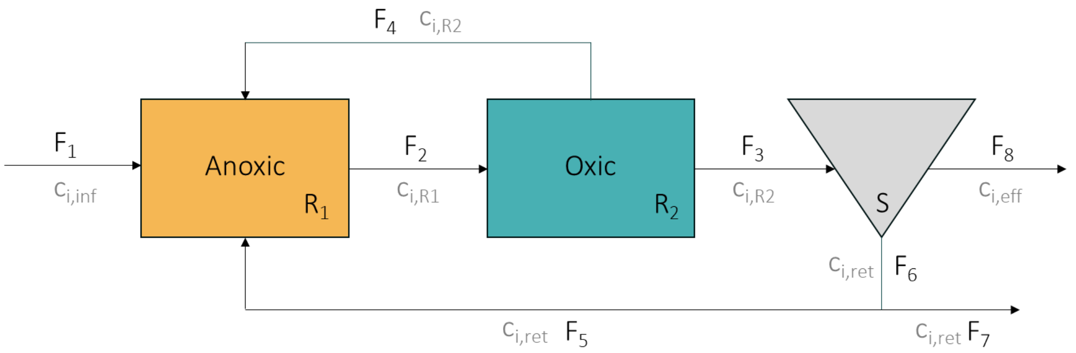

**Figure S22. Simplified flowsheet of the biological nutrient removal process used in the mathematical model, comprising an anoxic ($R_1$) and oxic tank ($R_2$), and a settler (S).** The flowrates are represented by $F_i$ and the metabolite and biomass concentrations in each process unit and stream are represented by $c_i$.

During the biological nitrogen removal process, denitrification occurs in the anoxic tank and nitrification in the oxic tank. For simplification, the denitrification process was not modelled in detail. Instead, to represent the $NO_3^-$ and $NO_2^-$ reduction in the anoxic tank, the rates were assumed to be very high compared to the ones of AOB and NOB, within the limits of the Monod terms (Table S9). The AOB and NOB nitrification conversions in the oxic tank were described with Monod kinetics (eq. S3 and S4). The fluctuation of $\mu_{max}$ with the temperature was described with eq. S5. The volumetric consumption or production rates in the oxic tank were determined from the specific growth rates (Table S9). AOB perform nitrifier denitrification to compensate limiting amounts of oxygen [39], so the $N_2O$ production rate by AOB was defined as dependent on the nitrite and oxygen concentrations (Table S9) [39,40]. Based on the $N_2O/NH_4^+$ emission factors found in an AOB enrichment by Peng et al. [40] (2-10%), the maximum $N_2O$ yield on biomass was assumed to be 2% of the ammonium yield (Table S10). The $O_2$ inhibition constant was assumed to be 1 $g \cdot m^{-3}$, according to Peng et al. [40].

**Table S9. Volumetric production and consumption rates of biomass and metabolites in the oxic and anoxic tanks.**

| Rate | Description | Formula | Units |
|---|---|---|---|
| **Anoxic – $R_1$** | | | |
| $r_{NO2,anx}$ | $NO_2^-$ reduction | $-100 \cdot \frac{c_{NO_2^-,R1}}{c_{NO_2^-,R1} + K_{NO_2^-}}$ | $gN \cdot m^{-3} \cdot d^{-1}$ |
| $r_{NO3,anx}$ | $NO_3^-$ reduction | $-100 \cdot \frac{c_{NO_3^-,R1}}{c_{NO_3^-,R1} + K_{NO_3^-}}$ | $gN \cdot m^{-3} \cdot d^{-1}$ |
| **Oxic – $R_2$** | | | |
| $r_{AOB}$ | AOB growth | $\mu_{max,AOB} \cdot c_{AOB} \cdot \frac{c_{O_2}}{c_{O_2} + K_{O_2}} \cdot \frac{c_{NH_4^+}}{c_{NH_4^+} + K_{NH_4^+}}$ | $gX \cdot m^{-3} \cdot d^{-1}$ |
| $r_{NOB}$ | NOB growth | $\mu_{max,NOB} \cdot c_{NOB} \cdot \frac{c_{O_2}}{c_{O_2} + K_{O_2}} \cdot \frac{c_{NO_2^-}}{c_{NO_2^-} + K_{NO_2^-}}$ | $gX \cdot m^{-3} \cdot d^{-1}$ |
| $r_{NH4,ox}$ | $NH_4^+$ oxidation by AOB | $-\frac{1}{Y_{AOB}} \cdot r_{AOB}$ | $gN \cdot m^{-3} \cdot d^{-1}$ |
| $r_{NO2,ox}$ | $NO_2^-$ production by AOB and oxidation by NOB | $\frac{1}{Y_{AOB}} \cdot r_{AOB} - \frac{1}{Y_{NOB}} \cdot r_{NOB}$ | $gN \cdot m^{-3} \cdot d^{-1}$ |
| $r_{NO3,ox}$ | $NO_3^-$ production by NOB | $\frac{1}{Y_{NOB}} \cdot r_{NOB}$ | $gN \cdot m^{-3} \cdot d^{-1}$ |
| $r_{N2O,ox}$ | $N_2O$ production by AOB | $Y_{N_2O} \cdot \mu_{max,AOB} \cdot c_{AOB} \cdot \frac{c_{NO_2^-}}{c_{NO_2^-} + K_{NO_2^-}} \cdot \frac{K_{I,O_2}}{c_{O_2} + K_{I,O_2}}$ | $gN \cdot m^{-3} \cdot d^{-1}$ |

**Table S10. Process and microbial parameters used in the mathematical model.**

| Parameter | Description | Value | Unit | Reference |
|---|---|---|---|---|
| **Process parameters** | | | | |
| SRT | Sludge retention time | 15 | d | This study |
| $V_1$ | Volume of $R_1$ (anoxic) | 11000 | $m^3$ | This study |
| $V_2$ | Volume of $R_2$ (oxic) | 5800 | $m^3$ | This study |
| $V_s$ | Volume of the settler | 18500 | $m^3$ | This study |
| $F_1$ | Influent flow rate | 26000 | $m^3 \cdot d^{-1}$ | This study |
| $F_4$ | Flow rate from the recycling from oxic to anoxic tank | 100000 | $m^3 \cdot d^{-1}$ | This study |
| $F_5$ | Flow rate of the return sludge | 26000 | $m^3 \cdot d^{-1}$ | This study |
| $c_{NH4,in}$ | Influent ammonium concentration | 50 | $gN \cdot m^{-3}$ | This study |
| **Stoichiometric and kinetic parameters** | | | | |
| $K_{O2,AOB}$ | $O_2$ half-saturation coefficient of AOB | 0.7 | $gO_2 \cdot m^{-3}$ | [41,42] |
| $K_{NH4,AOB}$ | $NH_4^+$ half-saturation coefficient of AOB | 1 | $gN \cdot m^{-3}$ | [43] |
| $K_{I,O2}$ | $O_2$ inhibition coefficient of nitrifier denitrification | 1 | $gO_2 \cdot m^{-3}$ | [40] |
| $\mu_{max,AOB}$ | Maximum specific growth rate of AOB | 0.20 | $d^{-1}$ | This study |
| $Y_{AOB}$ | AOB biomass yield on $NH_4^+$ | 0.13 | $gX \cdot gN^{-1}$ | Table S6 |
| $\theta_{AOB}$ | Arrhenius coefficient of AOB | 1.12 | - | This study |
| $Y_{N2O}$ | $N_2O$ yield on AOB biomass | 0.14 | $gN \cdot gX^{-1}$ | [40] |
| $K_{O2,NOB}$ | $O_2$ half-saturation coefficient of NOB | 0.1 | $gO_2 \cdot m^{-3}$ | [41,42] |
| $K_{NO2,NOB}$ | $NO_2^-$ half-saturation coefficient of NOB | 0.5 | $gN \cdot m^{-3}$ | [43] |
| $\mu_{max,NOB}$ | Maximum specific growth rate of NOB | 0.12 | $d^{-1}$ | This study |
| $Y_{NOB}$ | NOB biomass yield on $NH_4^+$ | 0.025 | $gX \cdot gN^{-1}$ | Table S6 |
| $\theta_{NOB}$ | Arrhenius coefficient of NOB | 1.10 | - | This study |

The unknown flow rates of the process were determined from the known flow rates with overall mass balances (Fig. S22, Table S11).

**Table S11. Overall mass balances to determine the unknown volumetric flow rates.**

| Flow rate | Description | Formula | Units |
|---|---|---|---|
| $F_2$ | Anoxic to oxic tank | $F_1 + F_4 + F_5$ | $m^3$ |
| $F_3$ | Oxic tank to settler | $F_2 - F_4$ | $m^3$ |
| $F_7$ | Purged sludge | $\frac{V_{R2} \cdot (c_{AOB,R2} + c_{NOB,R2})}{SRT \cdot (c_{AOB,ret} + c_{NOB,ret})}$ | $m^3$ |
| $F_6$ | Sludge effluent from settler | $F_5 + F_7$ | $m^3$ |
| $F_8$ | Water effluent from settler | $F_3 - F_6$ | $m^3$ |

Mass balances in each process unit described the dynamics in biomass and metabolite concentrations.

**Table S12. Mass balances (ordinary differential equations) of metabolites and biomass in each process unit.**

| Process unit | Ordinary differential equation | Compounds | Units |
|---|---|---|---|
| $R_1$ | $\frac{dc_{i,R1}}{dt} = \frac{c_{i,in} \cdot F_1 + c_{i,R2} \cdot F_4 + c_{i,ret} \cdot F_5 + c_{i,R1} \cdot F_2}{V_{R1}} + r_{i,R1}$ | Metabolites and biomass | $g \cdot m^{-3} \cdot d^{-1}$ |
| $R_2$ | $\frac{dc_{i,R2}}{dt} = \frac{c_{i,R1} \cdot F_2 + c_{i,R2} \cdot (F_3 + F_4)}{V_{R2}} + r_{i,R2}$ | Metabolites and biomass | $g \cdot m^{-3} \cdot d^{-1}$ |
| S | $\frac{dc_{i,ret}}{dt} = \frac{c_{i,R2} \cdot F_3 + c_{i,ret} \cdot F_6}{V_S}$ | Biomass | $g \cdot m^{-3} \cdot d^{-1}$ |
| | $\frac{dc_{i,ret}}{dt} = \frac{c_{i,R2} \cdot F_3 + c_{i,ret} \cdot (F_6 + F_8)}{V_S}$ | Metabolites | $g \cdot m^{-3} \cdot d^{-1}$ |

The ordinary differential equations were solved with the BDF method of the *solve_ivp* function of the SciPy v1.7.3 [44] package for a time span of 5 years. The oxygen concentration in the anoxic tank was set as 0. The seasonal fluctuations in the temperature and oxygen concentration in the oxic tank were replicated with opposite sinusoids as function of time:

$$\mathbf{c_{O_2}} \text{ or } \mathbf{T} = A \cdot \sin(2\pi \cdot f \cdot t + \varphi) + B \qquad \text{(eq. S7)}$$

**Table S13. Parameters for the sinusoids describing seasonal fluctuations in the oxygen concentration and the temperature.**

| Parameter | Description | Value for $c_{O2}$ sinusoid | Value for T sinusoid |
|---|---|---|---|
| Min | Minimum value | $0.5\ g \cdot m^{-3}$ | 10 °C |
| Max | Maximum value | $3\ g \cdot m^{-3}$ | 25 °C |
| A | Amplitude | $\frac{c_{O_2,max} - c_{O_2,min}}{2}$ | $\frac{T_{max} - T_{min}}{2}$ |
| f | Frequency | 1/365 | 1/365 |
| φ | Phase | $1.5\pi$ | $0.5\pi$ |
| B | Axis shift | $\frac{c_{O_2,max} + c_{O_2,min}}{2}$ | $\frac{T_{max} + T_{min}}{2}$ |

The model replicated the ecophysiological cascade, with the accumulation of $NH_4^+$, $NO_2^-$ and $N_2O$ and the increase in AOB/NOB ratio d(Fig. S22).

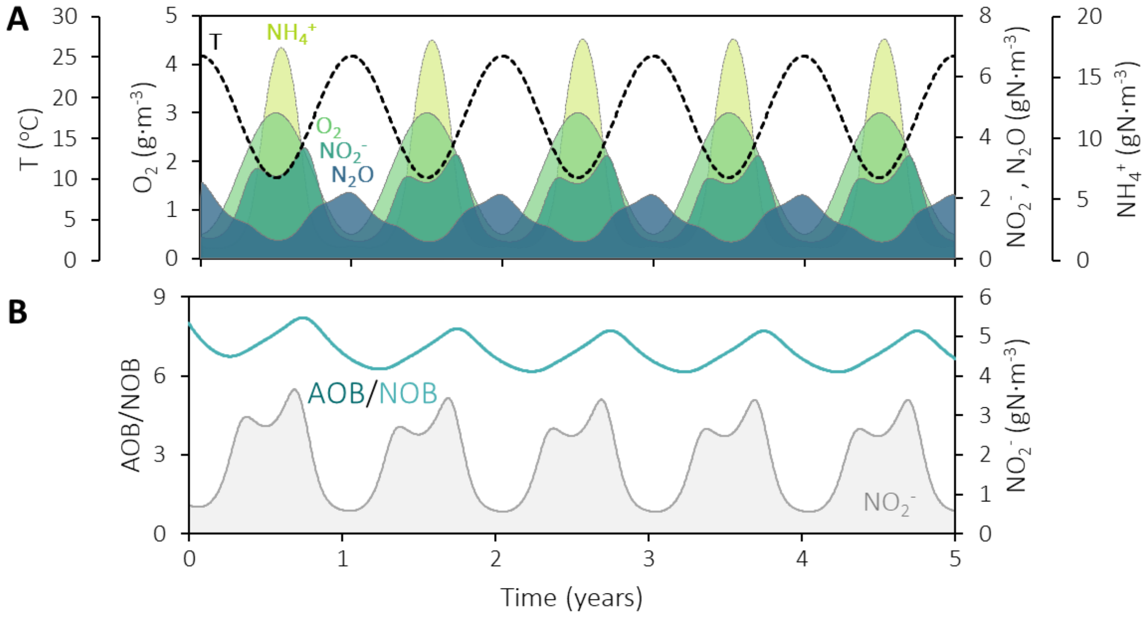

**Figure S23. Mathematical model simulating the seasonal nitrifiers and nitrogen metabolites dynamics.** In the model, the ecophysiological cascade was triggered with decreasing temperature and increasing dissolved $O_2$, as hypothesized from the full-scale observations. **(A)** Temperature, and ammonium, dissolved oxygen, nitrite and nitrous oxide concentrations in the oxic tank. **(B)** Ratio of the concentrations of AOB and NOB in the oxic tank, alongside the nitrite concentration also represented in A.

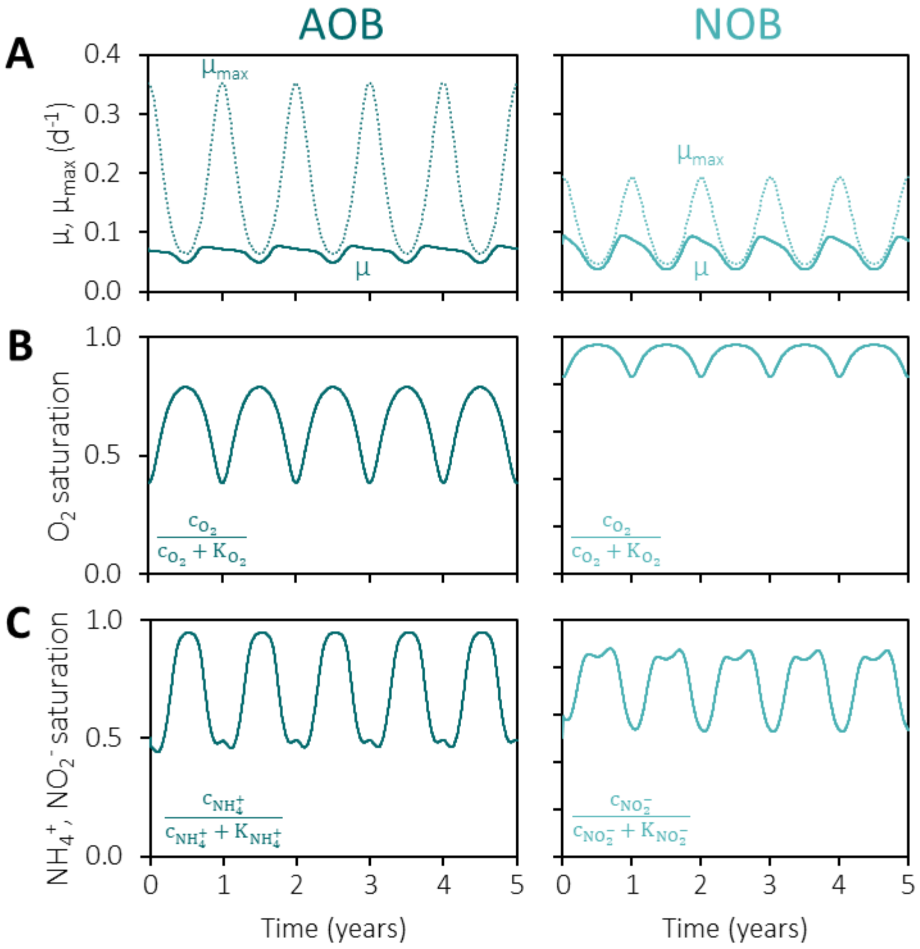

**Figure S24. Kinetic contributions of each element of the growth equations to the AOB and NOB growth rates (eq. S3-4) during temperature and dissolved oxygen fluctuations in the mathematical model represented in Fig. S23. (A)** Fluctuation of the maximum ($\mu_{max}$) and actual specific growth rates ($\mu$). While the $\mu_{max}$ of AOB decreases relatively more than the $\mu_{max}$ of NOB, the value for NOB achieves a much lower value, so the negative effect of temperature decrease harms NOB more than AOB. **(B)** Fluctuation in the dissolved oxygen saturation fraction. The relative increase in $\mu_{AOB}$ is much higher than in $\mu_{NOB}$, so the $O_2$ increase benefits AOB more than NOB. **(C)** Fluctuation in the ammonium (AOB) and nitrite (NOB) saturation fractions.

**Python script with the mathematical model replicating the seasonal nitrite accumulation**

```
import numpy as np
import scipy.integrate as spi
import matplotlib.pyplot as plt
from math import pi
from math import sin

# Assume a setup with an anoxic (R1) and aerobic reactor with fixed O2 concentration (R2),
# with a recycling stream from oxic to anoxic and the return sludge stream
# Implemented a temperature decrease with a simultaneous O2 increase
# AOB and NOB are competing for O2

#%% ----------------------- Model parameters -----------------------

# Process parameters
SRT    = 15            # days
V1     = 11000         # Volume reactor 1 (m3) - anoxic
V2     = 5800          # Volume reactor 2 (m3) - oxic
Vs     = 18500         # Volume settler (m3)
F1     = 26000         # Influent (m3/d)
F4     = 100000        # Recycling from oxic to anoxic (m3/d)
F5     = 30000         # Return sludge (m3/d)
F2     = F1 + F4 + F5  # R1 to R2
F3     = F2 - F4       # Effluent R2

# Stoichiometric and kinetic parameters
Ko2_aob   = 0.7    # O2   half-saturation coefficient AOB (g/m3)
Ko2_nob   = 0.1    # O2   half-saturation coefficient NOB (g/m3)
Knh4      = 1      # NH4+ half-saturation coefficient AOB (g/m3)
Kno2      = 0.5    # NO2- half-saturation coefficient NOB (g/m3)
Ki_o2     = 1      # O2   inhibition half-saturation coefficient for Nit Den (g/m3)
mum_aob   = 0.20   # mu_max of AOB (d-1)
mum_nob   = 0.12   # mu_max of NOB (d-1
Yaob      = 0.13   # AOB yield on NH4+ (gX/gN)
Yn2o      = 0.14   # N2O yield on AOB biomass (gN/gX)
Ynob      = 0.025  # NOB yield on NO2- (gX/gN)
theta_aob = 1.12   # Temperature coefficient AOB
theta_nob = 1.10   # Temperature coefficient NOB

# Inffluent concentrations
caob_in = 0            # gX/m3
cnob_in = 0            # gX/m3
cnh4_in = 50           # gN/m3
cno2_in = 0            # gN/m3
cno3_in = 0            # gN/m3
cn2o_in = 0            # gN/m3

#%% ----------------- Initial conditions for the ODE solver ---------------

# Initial concentrations - R1 anoxic tank
caob_r1_0 = 400        # gX/m3
cnob_r1_0 = 50         # gX/m3
cnh4_r1_0 = 1          # gN/m3
cno2_r1_0 = 0.5        # gN/m3
cno3_r1_0 = 5          # gN/m3
cn2o_r1_0 = 2          # gN/m3

# Initial concentrations - R2 oxic tank
```

```
caob_r2_0 = caob_r1_0   # gX/m3
cnob_r2_0 = cnob_r1_0   # gX/m3
cnh4_r2_0 = cnh4_r1_0   # gN/m3
cno2_r2_0 = cno2_r1_0   # gN/m3
cno3_r2_0 = cno3_r1_0   # gN/m3
cn2o_r2_0 = cn2o_r1_0   # gN/m3

# Initial concentrations - Return sludge
caob_re_0 = caob_r1_0*2 # gX/m3
cnob_re_0 = cnob_r1_0*2 # gX/m3
cnh4_re_0 = cnh4_r1_0   # gN/m3
cno2_re_0 = cno2_r1_0   # gN/m3
cno3_re_0 = cno3_r1_0   # gN/m3
cn2o_re_0 = cn2o_r1_0   # gN/m3

# Initial conditions array
c0 = [caob_r1_0, cnob_r1_0, cnh4_r1_0, cno2_r1_0, cno3_r1_0, cn2o_r1_0,
      caob_r2_0, cnob_r2_0, cnh4_r2_0, cno2_r2_0, cno3_r2_0, cn2o_r2_0,
      caob_re_0, cnob_re_0, cnh4_re_0, cno2_re_0, cno3_re_0, cn2o_re_0]

# Simulate for 5 years
t0, tf = 0, 5*365                      # Start and end time for the solver (d)
tspan  = np.linspace(t0, tf, tf+1)     # Time steps for which we want the solution

#%% --------------------- Define functions ----------------------

# Reproduce the seasonal O2 and T profiles:
# First half year: Increase O2 and decrease T
# Last half year:  Decrease O2 and increase T

# Define a function that returns the O2 concentrations for a certain time point
def co2_function(t):

    co2_min = 0.5   # g/m3
    co2_max = 3     # g/m3

    # Sine wave parameters
    A   = (co2_max - co2_min)/2   # amplitude
    f   = 1/365                   # frequency
    phi = 1.5*pi                  # phase
    B   = (co2_min + co2_max)/2   # shift from axis

    co2 = A*sin(2*pi*f*t + phi) + B

    return co2

# Define a function that returns the T for a certain time point
def T_function(t):

    T_min = 10    # oC
    T_max = 25    # oC

    # Sine wave parameters
    A   = (T_max - T_min)/2   # amplitude
    f   = 1/365               # frequency
    phi = 0.5*pi              # phase
    B   = (T_min + T_max)/2   # shift from axis
```

```
    T = A*sin(2*pi*f*t + phi) + B

    return T

# Define a function that returns the conversion rates based on the concentrations
def rates(caob, cnob, cnh4, cno2, cno3, cn2o, co2, T):

    # Growth rate at temperature T
    mum_aob_T = mum_aob * theta_aob ** (T - 20)
    mum_nob_T = mum_nob * theta_nob ** (T - 20)

    # Rate equations [g/m3/d]
    raob = mum_aob_T * caob * co2/(co2 + Ko2_aob) * cnh4/(cnh4 + Knh4)
    rnob = mum_nob_T * cnob * co2/(co2 + Ko2_nob) * cno2/(cno2 + Kno2)
    rnh4 = -1/Yaob * raob
    rno2 =  1/Yaob * raob - 1/Ynob * rnob
    rno3 =  1/Ynob * rnob

    # Hypothetical equation to represent N2O production through nitrifier denitrification
    rn2o   = Yn2o * mum_aob_T * caob * cno2/(cno2 + Kno2) * Ki_o2/(co2 + Ki_o2)

    return [raob, rnob, rnh4, rno2, rno3, rn2o]

# Define the ode system
def ode_cstr(t, c):

    # Concentrations inside the reactors [g/m3]
    caob_r1, cnob_r1, cnh4_r1, cno2_r1, cno3_r1, cn2o_r1, caob_r2, cnob_r2, cnh4_r2,
cno2_r2, cno3_r2, cn2o_r2, caob_re, cnob_re, cnh4_re, cno2_re, cno3_re, cn2o_re = c

    co2_r1 = 0                  # Anoxic reactor
    co2_r2 = co2_function(t)    # Oxic reactor
    T      = T_function(t)      # Temperature

    # Calculate total biomass concentration in each reactor: AOB + NOB
    cx_r2 = caob_r1 + cnob_r1
    cx_re = caob_re + cnob_re

    # Calculate the flows that maintain a constant SRT
    F7  = V2*cx_r2 / SRT / cx_re                    # Purged flow
    F6  = F5 + F7                                   # sludge effluent of settler
    F8  = F3 - F6                                   # water effluent of settler

    # Define the rate equations [g/m3/d] - R1 anoxic
    raob_r1, rnob_r1, rnh4_r1, rno2_r1, rno3_r1, rn2o_r1 = rates(caob_r1, cnob_r1, cnh4_r1,
cno2_r1, cno3_r1, cn2o_r1, co2_r1, T)
    rno2_r1 = - 100 * cno2_r1/(cno2_r1 + Kno2)  # Make it very high to assume complete
denitrification of NO2
    rno3_r1 = - 100 * cno3_r1/(cno3_r1 + Kno2)  # Make it very high to assume complete
denitrification of NO3

    # Define the rate equations [g/m3/d] - R2 oxic
    raob_r2, rnob_r2, rnh4_r2, rno2_r2, rno3_r2, rn2o_r2 = rates(caob_r2, cnob_r2, cnh4_r2,
cno2_r2, cno3_r2, cn2o_r2, co2_r2, T)

    # Define the ODEs [g/m3/d] - reactor 1 - anoxic
    dcaobdt_r1 = ( caob_in*F1 + caob_r2*F4 + caob_re*F5 - caob_r1*F2 ) / V1 + raob_r1
    dcnobdt_r1 = ( cnob_in*F1 + cnob_r2*F4 + cnob_re*F5 - cnob_r1*F2 ) / V1 + rnob_r1
```

```
    dcnh4dt_r1 = ( cnh4_in*F1 + cnh4_r2*F4 + cnh4_re*F5 - cnh4_r1*F2 ) / V1 + rnh4_r1
    dcno2dt_r1 = ( cno2_in*F1 + cno2_r2*F4 + cno2_re*F5 - cno2_r1*F2 ) / V1 + rno2_r1
    dcno3dt_r1 = ( cno3_in*F1 + cno3_r2*F4 + cno3_re*F5 - cno3_r1*F2 ) / V1 + rno3_r1
    dcn2odt_r1 = ( cn2o_in*F1 + cn2o_r2*F4 + cn2o_re*F5 - cn2o_r1*F2 ) / V1 + rn2o_r1

    # Define the ODEs [g/m3/d] - reactor 2 - oxic
    dcaobdt_r2 = ( caob_r1*F2 - caob_r2 * (F3 + F4) ) / V2 + raob_r2
    dcnobdt_r2 = ( cnob_r1*F2 - cnob_r2 * (F3 + F4) ) / V2 + rnob_r2
    dcnh4dt_r2 = ( cnh4_r1*F2 - cnh4_r2 * (F3 + F4) ) / V2 + rnh4_r2
    dcno2dt_r2 = ( cno2_r1*F2 - cno2_r2 * (F3 + F4) ) / V2 + rno2_r2
    dcno3dt_r2 = ( cno3_r1*F2 - cno3_r2 * (F3 + F4) ) / V2 + rno3_r2
    dcn2odt_r2 = ( cn2o_r1*F2 - cn2o_r2 * (F3 + F4) ) / V2 + rn2o_r2

    # Define the ODEs [g/m3/d] - settler
    dcaobdt_re = ( caob_r2*F3 - caob_re *  F6       ) / Vs
    dcnobdt_re = ( cnob_r2*F3 - cnob_re *  F6       ) / Vs
    dcnh4dt_re = ( cnh4_r2*F3 - cnh4_re * (F6 + F8) ) / Vs
    dcno2dt_re = ( cno2_r2*F3 - cno2_re * (F6 + F8) ) / Vs
    dcno3dt_re = ( cno3_r2*F3 - cno3_re * (F6 + F8) ) / Vs
    dcn2odt_re = ( cn2o_r2*F3 - cn2o_re * (F6 + F8) ) / Vs

    return [dcaobdt_r1, dcnobdt_r1, dcnh4dt_r1, dcno2dt_r1, dcno3dt_r1, dcn2odt_r1,
dcaobdt_r2, dcnobdt_r2, dcnh4dt_r2, dcno2dt_r2, dcno3dt_r2, dcn2odt_r2, dcaobdt_re,
dcnobdt_re, dcnh4dt_re, dcno2dt_re, dcno3dt_re, dcn2odt_re]

#%% ---------------- Solve the system of ODEs -----------------

sol = spi.solve_ivp(ode_cstr, [t0, tf], c0, t_eval = tspan, method = 'BDF')

# Retrieve the solutions for the timepoints specified in tspan
t       = sol.t
caob_r1, cnob_r1, cnh4_r1, cno2_r1, cno3_r1, cn2o_r1, caob_r2, cnob_r2, cnh4_r2, cno2_r2,
cno3_r2, cn2o_r2, caob_re, cnob_re, cnh4_re, cno2_re, cno3_re, cn2o_re = sol.y[0:18,:]

# Re-define co2 and T to calculate rates and plot the results
co2 = np.zeros(len(t))
T   = np.zeros(len(t))
for i in np.arange(len(t)):
    co2[i] = co2_function(t[i])
    T[i]   = T_function(t[i])

# Calculate rates to plot them
raob_r2, rnob_r2, rnh4_r2, rno2_r2, rno3_r2, rn2o_r2 = rates(caob_r2, cnob_r2, cnh4_r2,
cno2_r2, cno3_r2, cn2o_r2, co2, T)

#%% -------------- Plot the results ----------------

# Create a figure
plt.close("all")
plt.figure(figsize = (6, 8))

# colours for the plots
col_T   = '#B8B8B8'
col_o2  = '#cce762'
col_nh4 = '#83D387'
col_no2 = '#3DA38A'
col_n2o = '#376C8D'

# Plot the solutions
```

```
# subplot 1 - temperature
ax = plt.subplot(3, 1, 1)
ax.plot(t, T, c = col_T)
plt.legend(['T'])
plt.xlabel('Time (d)')
plt.ylabel('T ($^o$C)')
plt.ylim([0, 30])

# subplot 2 – NH4 and o2 on the left axis and NO2 and N2O on the right axis
ax = plt.subplot(3, 1, 2)
ax.plot(t, co2*5,   c = col_o2)
ax.plot(t, cnh4_r2, c = col_nh4)
plt.legend(['$c_{O2}$ *5', '$c_{NH4}$'], loc = "upper left")
plt.xlabel('Time (d)')
plt.ylabel('$c_{O2}$, $c_{NH4}$ (g/$m^3$)')
plt.ylim([0, 20])

ax2 = ax.twinx()
ax2.plot(t, cno2_r2, c = col_no2)
ax2.plot(t, cn2o_r2, c = col_n2o)
plt.legend(['$c_{NO2}$', '$c_{N2O}$'], loc = "upper right")
plt.ylabel('$c_{NO2}$, $c_{N2O}$ (g/$m^3$)')
plt.ylim([0, 6])

# subplot 3 – ratio AOB/NOB
ax = plt.subplot(3, 1, 3)
ax.plot(t, caob_r2/cnob_r2)
plt.legend(['AOB/NOB'])
plt.xlabel('Time (d)')
plt.ylabel('Ratio')
plt.ylim([0, 10])

plt.tight_layout()
```

### 13. Calculation of the maximum $N_2O$ activities

#### Determination of the $N_2O$ mass transfer coefficient (kLa)

The $N_2O$ volumetric mass transfer coefficient ($k_La$) was determined to calculate the transfer rate during the $N_2O$ batches. The $k_La$ was determined under the same conditions as the activity tests (750 rpm stirring, no gas flow, 20 ºC), with water instead of biomass. The $k_La$ was obtained by taking the slope of the linearized integrated mass transfer equation fitted to the dissolved $N_2O$ concentration profile over time.

$$C_{N_2O} = C^*_{N_2O} \cdot (1 - e^{-k_La \cdot t}) \quad \text{(eq. S8)}$$

with $C_{N2O}^*$ the solubility of $N_2O$ at 20 °C. The obtained $k_La$ was 5 $h^{-1}$.

#### Python script to calculate the maximum $N_2O$ activities

```
' ------ Purpose of the code ------ '
# Determine the maximum N2O consumption rate (rN2O) during a series of batch tests.
# This is done by fitting the model of N2O dynamics (considering reaction and gas-liquid
transfer) to the experimental data.

import pandas
from scipy.integrate import solve_ivp
from scipy.optimize import minimize
import numpy as np

#%%   ----------- Import the raw data -----------

# Load the data from all batches
# column 1 is the time in min
# the remaining 29 columns are concentration profiles from each batch
data_import = pandas.read_excel('conc_for_python_script_N2O.xlsx')

# Separate the time and concentration in two different variables
time_exp = np.arange(len(data_import))       # 0-74 min
conc_exp = data_import.iloc[:,1:]/28         # Convert to mM

# Constants
Kh  = 27.05      # Henry constant (mM/atm)
p   = 1          # Pressure (atm)
R   = 0.00008206 # Ideal gas constant (L.atm/K/mmol)
T   = 20+273.15  # Temperature (K)
kLa = 5/60       # Experimentally determined kLa (converted from 1/h to 1/min)
V   = 2          # Liquid volume (L)
Vg  = 1          # Headspace volume (L)

# Make an array with zeros to save solutions
rates = np.zeros((len(conc_exp.columns),1))

#%%   ----------- Define functions -----------

' -------- System of odes -------- '

# We assume rN2O = rN2O_max because concentrations are very high
# Arguments of the ODE function: time, concentrations and the unknown N2O rate

def ode_n2o(t, x, rN2O):

    # The variables are N2O concentration in liquid and gas
    Sn2o, Sn2og = x              # mM
```

```
    # Calculate N2O solubility and gas-liquid transfer rate based on the
    # current liquid and gas concentrations
    Sn2os = Kh*Sn2og*R*T/p       # mM
    rN2Ot = kLa*(Sn2o - Sn2os)   # mM/min

    # The ODEs are the mass balances of N2O in the liquid and gas
    dsdt  = rN2O - rN2Ot         # mM/min - Liquid
    dsgdt = rN2Ot * V/Vg         # mM/min - Gas

    return [dsdt, dsgdt]

'--- Minimization of the SSE between predicted and experimental conc N2O ---'

# Arguments of the objective function:
# Parameter to be determined: rN2O
# Initial guess: x0
# Experimental time points: t_exp
# Experimentally measured concentrations: c_exp

def n2o_SSE(rN2O, x0, t_exp, c_exp):

    # Time interval for the solver
    tspan = [t_exp[0], t_exp[-1]]

    # Solve the system of ODEs for each iteration of the minimizer
    # ode_n2o:   function containing the ODEs
    # x0:        values of the variables (Sn2o & Sn2og) at time 0
    # t_eval:    timepoints for which we want an output
    # method:    RK45 for non-stiff systems
    # args:      pass the parameter as argument to the ode_n2o function
    sol = solve_ivp(ode_n2o, tspan, x0, t_eval = t_exp, method = 'RK45', args = (rN2O),
rtol = 1e-12)

    # Retrieve the solution - N2O concentration in the liquid (mM)
    Sn2o  = sol.y[0, :]

    # Calculate the individual sum of squared errors between calculated and
    # experimental N2O concentrations in the liquid
    SSE = sum((Sn2o - c_exp)**2)

    return SSE

' ------- Take each batch dataset in turns -------- '

# Run loops through each batch test
for batch in np.arange(len(conc_exp.columns)):

    # Remove NaN rows
    c_exp = conc_exp.values[:,batch]    # select the right column from the experimental
data
    t_exp = time_exp[~np.isnan(c_exp)]  # Remove the time points for which the
concentration was NaN
    c_exp = c_exp[~np.isnan(c_exp)]     # Remove the NaN elements

    # Set time points to 0 (in case the first point was removed above)
    t_exp = t_exp - t_exp[0]     # min
```

```
    # Concentrations at the beginning of the batch (Sn2o and Sn2og)
    Sn2o_0  = c_exp[0]           # N2O in the liquid    (mM) - experimental
    Sn2og_0 = Sn2o_0*1/Kh/R/T    # N2O in the headspace (mM) - calculated
    x0 = [Sn2o_0, Sn2og_0]

    # Initial guess for the rate
    rN2O_0 = -0.01        # mM/min

    # Batch 15 was very slow so we need a different initial guess for convergence
    if batch == 15:
        rN2O_0 = -0.001   # mM/min

    # Perform the minimization on the SSE function
    # n2o_SSE:   objective function for the minimization
    # rN2O_0:    initial guess for the rate
    # method:    minimization method
    # args:      pass the starting and experimental data to the minimization function
    res = minimize(n2o_SSE, rN2O_0, method = 'Nelder-Mead', tol = 1e-12, args = (x0, t_exp,
c_exp))

    # Retrieve the results
    rN2O = res.x    # mM/min

    # Save results in arrays and convert units from mM/L/min to mgN/L/h
    rates[batch] = abs(rN2O*28*60)

# Save the data
rates_save = pandas.DataFrame(rates[:,0]) # mgN/L/h
rates_save.to_csv('rates_N2O.csv')
```
